# Environmental and physiological factors affecting high-throughput measurements of bacterial growth

**DOI:** 10.1101/2020.06.16.156182

**Authors:** Esha Atolia, Spencer Cesar, Heidi A. Arjes, Manohary Rajendram, Handuo Shi, Benjamin D. Knapp, Somya Khare, Andrés Aranda-Díaz, Richard E. Lenski, Kerwyn Casey Huang

## Abstract

Bacterial growth in nutrient-rich and starvation conditions is intrinsically tied to the environmental history and physiological state of the population. While high-throughput technologies have enabled rapid analyses of mutant libraries, technical and biological challenges complicate data collection and interpretation. Here, we present a framework for the execution and analysis of growth measurements with improved accuracy over standard approaches. Using this framework, we demonstrate key biological insights that emerge from consideration of culturing conditions and history. We determined that quantification of the background absorbance in each well of a multi-well plate is critical for accurate measurements of maximal growth rate. Using mathematical modeling, we demonstrated that maximal growth rate is dependent on initial cell density, which distorts comparisons across strains with variable lag properties. We established a multiple-passage protocol that alleviates the substantial effects of glycerol on growth in carbon-poor media, and we tracked growth rate-mediated fitness increases observed during a long-term evolution of *Escherichia coli* in low glucose concentrations. Finally, we showed that growth of *Bacillus subtilis* in the presence of glycerol induces a long lag in the next passage due to inhibition of a large fraction of the population. Transposon mutagenesis linked this phenotype to the incorporation of glycerol into lipoteichoic acids, revealing a new role for these envelope components in resuming growth after starvation. Together, our investigations underscore the complex physiology of bacteria during bulk passaging and the importance of robust strategies to understand and quantify growth.

**Abstract Importance:** How starved bacteria adapt to and multiply in replete nutrient conditions is intimately linked to their history of previous growth, their physiological state, and the surrounding environment. While automated equipment has enabled high-throughput growth measurements, data interpretation and knowledge gaps regarding the determinants of growth kinetics complicate comparisons between strains. Here, we present a framework for growth measurements that improves accuracy and attenuates the effects of growth history. We determined that background absorbance quantification and multiple passaging cycles allows for accurate growth-rate measurements even in carbon-poor media, which we used to reveal growth-rate increases during long-term laboratory evolution of *Escherichia coli*. Using mathematical modeling, we showed that maximum growth rate depends on initial cell density. Finally, we demonstrated that growth of *Bacillus subtilis* with glycerol inhibits the future growth of most of the population, due to lipoteichoic-acid synthesis. These studies highlight the challenges of accurate quantification of bacterial growth behaviors.

## Introduction

Precise growth measurements are fundamental to our understanding of bacterial physiology and its regulation. While some bacterial species are among the fastest-growing organisms on the planet, others grow imperceptibly slowly, with doubling times ranging from ∼7 minutes [1] to thousands of years [2]. Although the need for rapid growth may drive selection in some cases, many bacteria live in complex natural environments that are often stressful and nutrient limited [3]. In many environments, such as the mammalian gut, leaf litter in soil, and whale falls in the ocean, food is provided only periodically, and hence bacteria experience cycles of feast and famine. Thus, the transition from starvation to rapid growth can also act as an important selective pressure in evolution [4-6].

A classical batch laboratory assay that encompasses all phases of growth involves the initial overnight growth of a liquid culture, from either a frozen stock or a colony, which is then used to inoculate fresh medium for optical-density (OD) measurements in a plate reader or spectrophotometer over time [7] (Fig. 1A). Although there are exceptions when OD does not track viable cell number [8, 9], OD is widely used as a proxy for the density of cells in a culture [10, 11]. While traversing such a growth curve, a cell population initially takes some time to accelerate in growth, experiences a period of rapid growth, and then decelerates as nutrients are consumed, waste products accumulate, or both [7]. While there can be substantial variability in the shape of the growth curve, many species qualitatively exhibit a sigmoidal shape that can be characterized by three parameters: (1) the maximal growth rate, *μ*_max_, which is the largest slope of the natural log of the OD over time, (2) the lag time required to accelerate growth from stationary phase (*T*_lag_), and (3) the maximum or final OD (*A*_max_); some cultures also exhibit a death phase [7]. These parameters are typically extracted from a growth curve via fitting or direct calculation [12-21]. Long-term evolution experiments (LTEEs) have demonstrated that all three can be under selection [4, 22], underscoring the importance of their accurate quantification. However, we have little understanding of how the technical aspects of model fitting and the methodological aspects of inoculation affect such quantifications.

**Figure 1:**
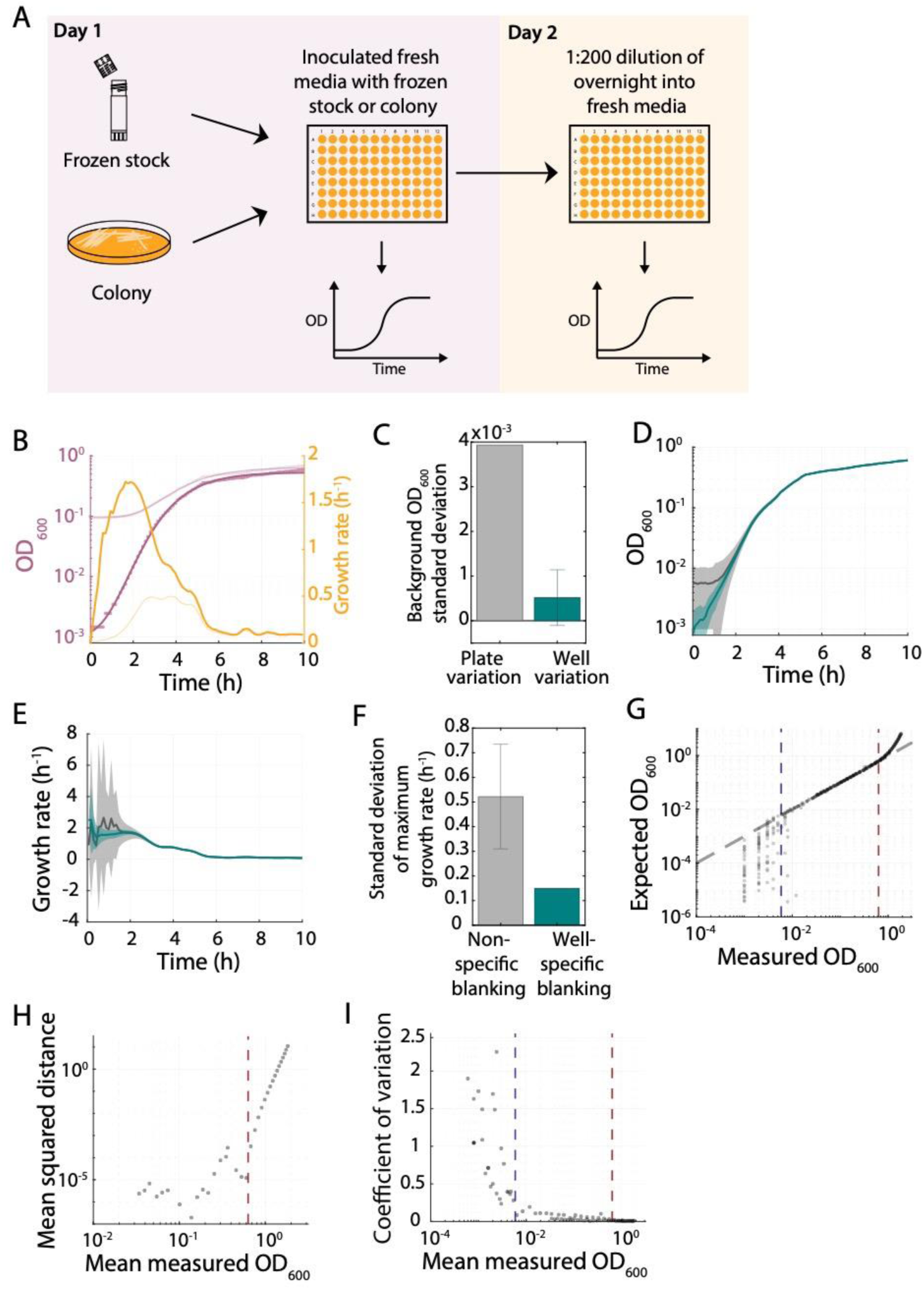
Well-specific blanking and establishment of the spectrophotometer limit of detection are critical for accurate measurements of population growth rate. A) Schematic of a classical experimental setup to measure bacterial growth. B) Raw absorbance values from a typical *E. coli* MG1655 growth curve (light purple curve) increased starting from just above the background absorbance of the well plus media (∼0.08). Subtracting the background OD resulted in absorbance values (dark purple circles) that indicate substantially different growth kinetics, which are well fit by a Gompertz function (dark purple curve). With background subtraction, the maximum growth rate was 3.7-fold higher and occurs ∼60 min earlier (dark yellow curve) than without subtraction (light yellow curve). C) The variation in the OD of a cell-free well containing only medium (gray) is much higher across a 96-well plate than the mean variation across multiple readings from a single well (teal). D) The variation in *E. coli* growth curves across a 96-well plate when blanking with the cell-free absorbance of each well (teal) was much lower than when blanking with the background absorbance of a random well (black). Shaded regions represent ±1 standard deviation. E) Instantaneous growth rates computed from growth curves in (B) were much less variable for well-specific blank subtraction (teal) than for blanking with a random well (black). F) Well-specific blank subtraction (teal) dramatically decreases the standard deviation in the estimate of maximum growth rate compared with subtraction of the blank from a randomly selected well in a 96-well plate (gray). G) A dilution series from a culture with a known OD can be used to calibrate OD readings in order to establish the range of linearity and the limit of detection of a plate reader. Blue and red dotted lines represent the linear range of detection; even above the linear range the variability is low. H) The mean squared distance from the line *x* = *y* in (D) increased sharply above the red dotted line, justifying its definition as the upper limit of the range of linearity. I) The coefficient of variation (CV, mean/standard deviation) of OD values in (D) increased sharply below the blue dotted line, justifying its definition as the lower limit of detection. Given the low CV above the linear range (above the red dotted line), it is possible to accurately measure up to an actual OD∼3.5 through correction based on the calibration in (G). The red dotted line represents the upper limit of the linear range.

The creation of genome-wide knockout libraries [23, 24] has motivated high-throughput measurements of population growth—a commonly used proxy for fitness—in microtiter plates, such as the systems-biological characterization of essential gene knockdowns in *Bacillus subtilis* [5] and of *Escherichia coli* non-essential gene knockouts in liquid [25, 26] or embedded in a gel [26, 27]. As the development of advanced genetic tools simplifies the creation of strain libraries [28-30], it is critical to ensure that OD measurements in multi-well plates provide reliable estimates of population-level growth parameters.

Here, we provide a detailed analysis of the requirements for achieving robust, high-throughput measurements of fitness parameters. We quantified the importance of accurate background subtraction and established an assay of the sensitivity of the plate reader. We determined that measurements of maximal population growth rate and lag time are both sensitive to initial inoculation density, and we developed a mathematical model based on ordinary differential equations that accurately predicts this density dependence. We showed that completely removing residual glycerol from frozen stocks as a carbon source is critical for accurate measurements of *E. coli* growth in carbon-limited media, and we used such a strategy to reveal that evolution in a carbon-poor medium led to significant increases in the maximal growth rate of a population. We also established that the presence of glycerol during initial outgrowth from frozen stocks can have long-term impact on *B. subtilis* growth, causing long apparent lag times during subsequent culturing due to growth inhibition in a large majority of cells. Using transposon mutagenesis, we discovered that the increased lag time at least partially results from incorporation of glycerol during lipoteichoic acid synthesis. For large-scale, high-throughput experiments, inoculating a large number of cultures from colonies is often too cumbersome, and thus these considerations are vital. Together, these findings provide a framework for accurate quantification of growth parameters and a roadmap for identifying and controlling for physiological factors that impact growth.

## Results

### Accuracy of population density estimation from spectrophotometer absorbance readings is sensitive to the method of background subtraction

To monitor maximum growth rate and lag time (defined here as time to reach half-maximum growth rate) during a bacterial population’s exit from stationary phase, it is standard practice to dilute a stationary-phase culture sufficiently that the spent medium transferred with the cells is a small fraction of the solution relative to the fresh medium. For a 100- to 1000-fold dilution of a stationary-phase culture with OD ∼1 (typical for many species in high-nutrient conditions measured with our plate reader), the starting OD is ∼0.001-0.01. Thus, for species such as *E. coli* for which the maximum growth rate is achieved within 2-3 generations [31], the corresponding OD at the time of maximum growth rate can be low (≲0.01) (Fig. 1A), making it critical to ascertain whether OD can be accurately measured at low cell densities.

It is generally appreciated that correcting for the background absorbance improves the accuracy of growth measurements. However, the extent to which different background correction methods affect growth rate calculations has not been quantified. For a culture that is growing exponentially, the number of cells, *N*, grows over time as *N*(*t*) = *N*_0_2^*t*/*τ*^, where *N*_0_ is the number of cells at *t* = 0 and *τ* is the doubling time. Thus, the growth rate of such a culture can be defined as the constant

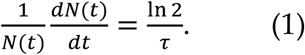

During outgrowth from stationary phase, when cells adapt their proteome to exploit the newly available nutrients [32], or after the cell density is sufficiently high that growth modifies the environment in a manner that impacts cellular physiology, the population does not grow exponentially. Nonetheless, analogous to exponential growth, we can define an instantaneous growth rate as

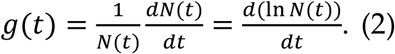

Assuming that the OD measured by a plate reader is linearly related to *N* (OD = *αN*, where *α* is a scaling factor relating cell number to OD) and measurements are taken at time points *t, t* + Δ*t, t* + 2Δ*t*, etc., the instantaneous growth rate at time *t* can be estimated as

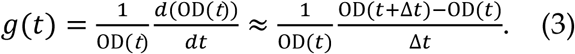

To illustrate the importance of background subtraction, consider a culture in which OD_raw_(*t*) = *αN*(*t*) + OD_bg_, where OD_bg_ is the background absorbance in the absence of cells; OD_bg_ is typically ∼0.1 (Fig. 1A). Without subtraction of OD_bg_, the computed growth rate would be 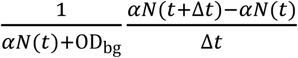 rather than 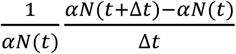; the estimate of growth rate would be incorrect by a factor of 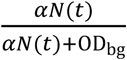 (Fig. 1A). Thus, any estimate of growth rate without background (blank) subtraction is highly underestimated when in the regime OD_bg_ ≳ *αN*(*t*), which is likely for the time at which the maximum growth rate of many bacterial species is first achieved. For similar reasons, using the first time point of a growth curve as a proxy for the background absorbance leads to overestimation of the maximum growth rate because the subtracted background was too large. Therefore, a method for correctly subtracting the background is crucial for accurate growth-rate estimates.

We measured growth curves of *E. coli* MG1655 and estimated the instantaneous growth rate over time (Methods), with and without blank subtraction. After subtracting the well blank, the maximal growth rate was 1.83 h^-1^ (doubling time of 22.7 min), which occurred at *t* = 1.52 h (Fig. 1B). Without blank subtraction, the maximal growth rate estimate was substantially lower (0.49 h^-1^) and the time at which this inaccurate estimate occurred was *t* = 3.0 h (Fig. 1B), illustrating the effects of omitting blank subtraction on lag time. To determine whether one empty well could be used as a general proxy for background absorbance, we quantified the absorbance of each well with medium before inoculating cells (Methods). Blank values varied by ∼0.004 across the plate, while a single well’s blank value fluctuated by <0.001 over time (Fig. 1C). Blanking with a randomly selected well from the plate led to a wide variation in blanked growth curves (Fig. 1D) and maximum growth rates (Fig. 1E) with a standard deviation in growth-rate estimate of 0.52 h^-1^ (Fig. 1F). Background subtraction with a well-specific blank led to substantially less variation in growth-rate measurements, with a standard deviation in growth-rate estimates for replicate cultures across the plate of 0.15 h^-1^ (Fig. 1F). Well-specific blanking also decreased the variability in lag measurements (Fig. S1A). The within-plate variability is sufficient to change the rank ordering of growth rates, which can lead to erroneous inferences. Thus, well-specific blanking is critical for accurate measurements of growth rate, because even comparisons in a single plate are confounded by within-plate variability.

### Sensitivity limit of the spectrophotometer also impacts the accuracy of growth-rate measurements

The ability to accurately measure changes in OD across a range of cell densities spanning several orders of magnitude is equally important for growth-rate calculations. Thus, we sought a general protocol for quantifying the limit of sensitivity and linear range of a given spectrophotometer. We inoculated serial dilutions of an overnight culture of *E. coli* MG1655 into fresh LB and measured the OD values. At low dilutions (OD>0.63), the absorbance measured by the plate reader was not linearly related to cell density (Fig. 1G,H). Nonetheless, high OD measurements could be converted to cell-density estimates with a measured calibration curve (Fig. 1G), because the measurement coefficient of variation (CV; standard deviation/mean) remained very low (Fig. 1I). To determine the precision of plate-reader measurements at high density, we converted the spread in growth-curve replicates at the same dilution into standard deviations of cell-density estimates, and found that the range of accurate measurement could be reliably extended up to the maximum tested expected OD of ∼3.5 (Fig. 1G), which is substantially higher than the typical corrected OD of ∼1 for a stationary phase *E. coli* culture grown in LB.

For OD values below ∼1, absorbances after well-specific blanking were linearly related to the dilution factor (Fig. 1G) and had a CV <0.2 (Fig. 1I) down to a blank-corrected OD of ∼0.006, well below the plate background OD_bg_. For larger dilutions (OD<0.006), the CV increased sharply (Fig. 1I), indicating that growth-rate measurements in our spectrophotometer are likely to be very noisy for OD <0.006. We conclude that any growth curve should be initialized with an inoculum such that the maximum growth rate is achieved above a blank-corrected value of 0.006. This strategy is likely effective for calibrating and determining the sensitivity limit of any spectrophotometer.

### Maximum growth rates can depend strongly on initial inoculation density

We previously found that the instantaneous population growth rate was strongly correlated with OD across a library of *B. subtilis* mutants, despite their widely varying lag times, suggesting that cell density plays a major role in determining the population’s growth rate [5]. Thus, to determine the optimal dilution for initializing growth curves, we systematically quantified how the inoculum cell density affects the trajectory of outgrowth from stationary phase. We diluted an overnight culture of *E. coli* MG1655 into fresh LB at ratios ranging from 1:12.5 to 1:6400 in a 96-well plate, and monitored the growth curves (Fig. 2A). To examine the relationship between OD and growth rate, we plotted each curve as growth rate versus log_10_(OD) (Fig. 2B). After correcting for the nonlinearity at high ODs (Fig. 1G), and subtracting the well-specific backgrounds, we found that the maximum growth rate achieved at low dilutions (e.g., 1:12.5) was lower than at larger dilutions (Fig. 2B). As expected, each curve started at a different initial OD with a growth rate near 0 (Fig. 2B). Growth rate then increased, and for large dilutions (1:6400) reached *μ*_max_ ≈ 2 h^-1^ (Fig. 2B). However, for lower dilutions, *μ*_max_ was substantially less than 2 h^-1^ and was attained at higher OD (Fig. 2B). In each case, after reaching *μ*_max_ growth rate declined approximately linearly as a function of log_10_(OD) along a common trajectory (Fig. 2B). Thus, before a population reaches its maximum growth rate, its growth rate trajectory is dependent on the initial cell density; thereafter, the growth curve follows a prescribed path independent of initial cell density.

**Figure 2:**
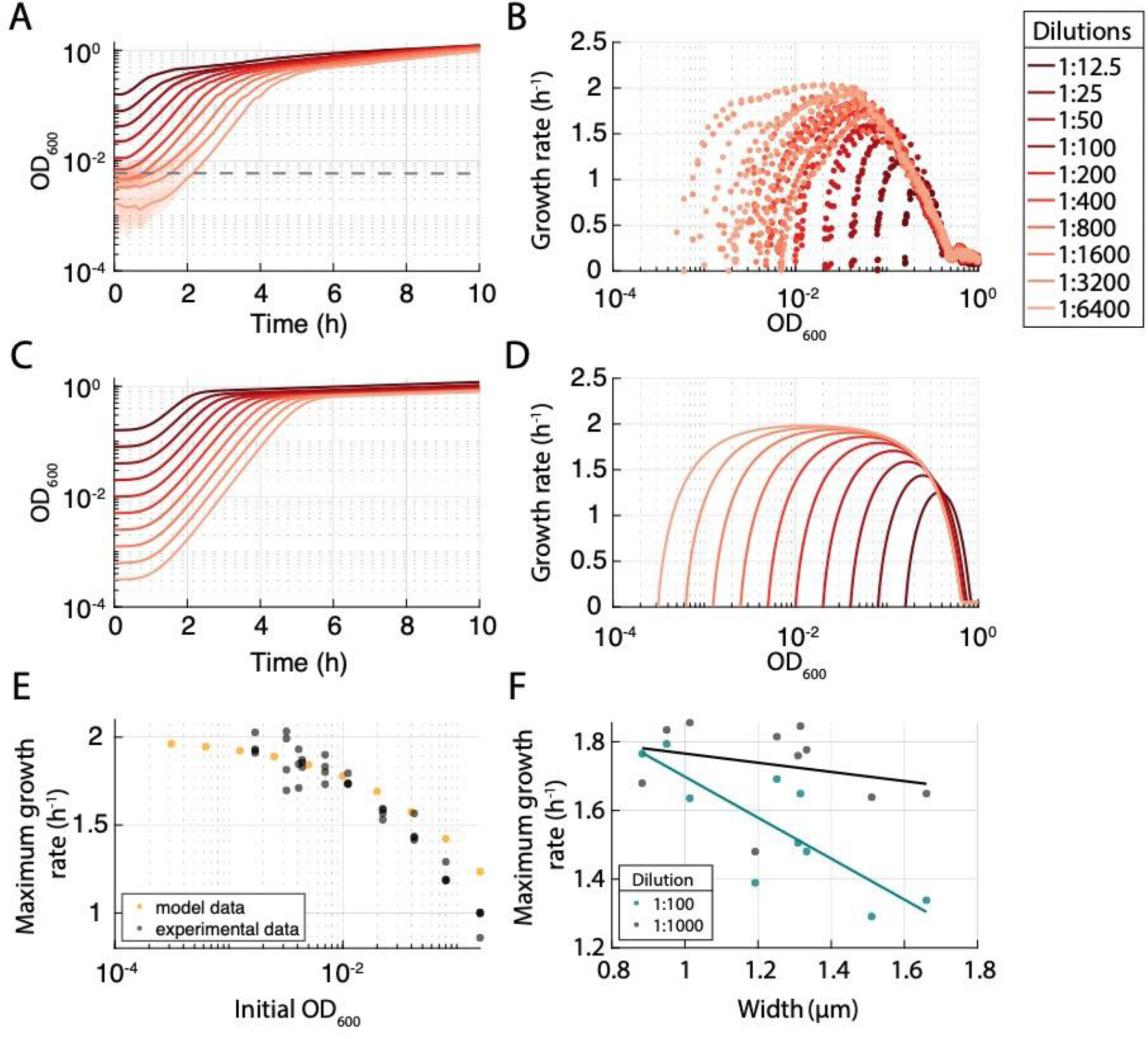
Growth rate is intrinsically linked to cell density due to nutrient depletion. A) Growth curves of a dilution series from a single overnight culture of *E. coli* MG1655 displayed distinct growth behaviors, with slower initial growth for lower dilutions (higher initial cell density). B) Instantaneous growth rate as a function of OD for the curves in (B) show that lower dilutions result in lower maximum growth rates. Curves followed a common, approximately linear downward trajectory after reaching maximum growth rate, indicating that the entry to stationary phase is less affected by initial OD than lag time or maximum growth rate. C) Nutrient depletion was sufficient to recapitulate experimental growth curves. Simulated growth curves for a model of growth based on nutrient depletion in Eqs. 4-6 were similar to experimental data in (B). D) Nutrient depletion was sufficient to recapitulate experimental relationship between OD and instantaneous growth rate. Instantaneous growth rate as a function of OD for the curves in (C) exhibited similar behaviors to the experimental data in (B). H) Nutrient depletion was sufficient to recapitulate experimental relationship between initial inoculum size and maximum growth rate. The maximum growth rates of the experimental (B) and simulated growth curves (D) exhibited a quantitatively similar decrease with increasing initial inoculum density. F) For a library of MreB mutants [34], the maximum growth rate computed from a growth curve initialized with a 1:100 dilution of an overnight culture (teal) decreased strongly with the mean cell width, while curves initialized from a 1:10,000 dilution (black) displayed higher, roughly constant maximum growth rates. The teal and black lines are least-squares linear fits to the data.

To interrogate whether factors such as nutrient depletion or waste accumulation cause this density dependence, we developed a minimal model of population growth dynamics during passage in liquid culture. We assume that cell density *C* grows with an instantaneous growth rate *μ* dictated by the physiological status of the cells and the external environment:

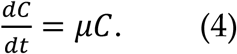

Nutrients are consumed by growing cells at a rate proportional to their growth rate:

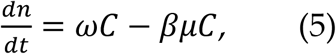

where *n* is nutrient concentration, ω represents production of nutrients by the cells, and *β* is the nutrient consumption rate. We assume that growth rate is a function of the nutrient concentration relative to a fixed nutrient concentration *K* [7]; to model the transition from stationary phase into log phase, we assume that *µ* is related to the highest possible maximal growth rate *μ*^*^ via a Gompertz relation [33]:

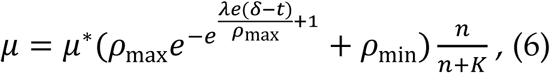

where *ρ*_min_ is the lowest growth rate attained at high nutrient concentration normalized to *µ*^*^, *ρ*_max_ = 1 − *ρ*_min_, *λ* governs how quickly *µ* increases, and *δ* is the time required to reach the maximum rate of growth-rate change. We used single-cell growth data to obtain estimates of *ρ*_max_, *ρ*_min_, *λ*, and *δ* (Fig. S1B). We found that the simulated growth curves were relatively insensitive to the precise functional form of the acceleration in growth during lag phase.

We simulated growth curves based on Eqs. (4-6), assuming that OD is proportional to *C*, with different initial densities *C*(*t* = 0) and *K* = 0.5 (where *n* = 1 is the maximum nutrient level), *μ*^*^ = 2 h^−1^, *β* = 0.8 h, *ρ*_max_ = 0.99, *ρ*_min_ = 0.01, *λ* = 0.8 h^−1^, and *δ* = 0.5 h. The kinetics of these growth curves (Fig. 2C) and the resulting relation between growth rate and OD (Fig. 2D) recapitulated our experimental findings reasonably well (Fig. 2B), including the roughly linear decrease in growth rate with OD after reaching *μ*_max_. Hence, nutrient depletion can largely explain the relations between *μ*_max_ and inoculation density (Fig. 2E), between lag and inoculation density (Fig. S1C), and more generally between growth rate and OD (Fig. 2D).

To illustrate the importance of these results, we examined the growth of a library of *E. coli* cell-size mutants [34]. After a 1:10,000 dilution, all mutants exhibited similar maximum growth rates (Fig. 2F). However, after a 1:200 dilution, maximum growth rate was negatively correlated with the average cell width of each mutant (Fig. 2F) [34]. This effect appears to reflect differences in outgrowth that prevented many of the mutants from attaining the higher maximum growth rate achievable at lower inoculation densities. These findings illustrate the importance of initiating growth curves with as low a cell density as possible without dropping below the plate reader’s limit of detection, in order to avoid the region of decreasing maximum growth rate at high cell densities (Fig. 2F) that can distort comparisons between strains.

### Growth in a carbon-poor medium is highly sensitive to glycerol levels

High-throughput methods often involve inoculation directly from a frozen stock rather than passaging through colonies, which could result in the transfer of variable amounts of glycerol, a cryoprotectant that ameliorates cell death during storage at low temperature [35, 36]. Thus, we sought to identify factors, such as glycerol, that affect the growth of cultures inoculated directly from frozen stock and then passaged multiple times. We hypothesized that glycerol would have persistent effects on growth, particularly in nutrient-poor media, because it can be utilized as a carbon source. Glycerol use that substantially increases the number of cells during the first passage would then perturb growth in later passages by changing the subsequent inoculation density (Fig. 2A). Such conditions are particularly relevant for strains generated by evolution experiments, which are often carried out in media with low carbon concentrations [37]. To test the effect of glycerol on growth, we measured growth curves across a range of glycerol concentrations (Fig. 3A-D). We inoculated 1 µL from a −80 °C freezer stock (previously grown in Davis Minimal medium with 25 g/L glucose, DM25) of *E. coli* REL606 [38], the ancestral strain for the multi-decade long-term evolution experiment (LTEE) carried out by Lenski and colleagues, into the evolution medium (DM25) supplemented with 0-10% (v/v) glycerol (Fig. 3A,B). The addition of glycerol mimics various levels of glycerol carryover from frozen stocks during inoculation. (However, we emphasize that the growth rate and competitive fitness assays performed by Lenski and collaborators prevent carryover of glycerol through repeated culturing in DM25, which acclimates the bacteria to the medium and other conditions of the LTEE prior to the fitness assays [39]). When we diluted these cultures 1:200, the resulting initial ODs substantially differed across glycerol concentrations, resulting in different growth kinetics with higher final OD values for cultures coming from higher glycerol concentrations on day 2 (Fig. 3C,E). After passaging the cultures a third time, the growth curves for the various glycerol concentrations were now quantitatively similar (Fig. 3D) with lower final ODs, as expected (Fig. 3E). Similar glycerol-dependent effects appeared when different amounts of frozen stock were inoculated into DM25 (Fig. S2A,B). Thus, accurate quantification of growth requires multiple passages to eliminate the effects of glycerol on growth.

**Figure 3:**
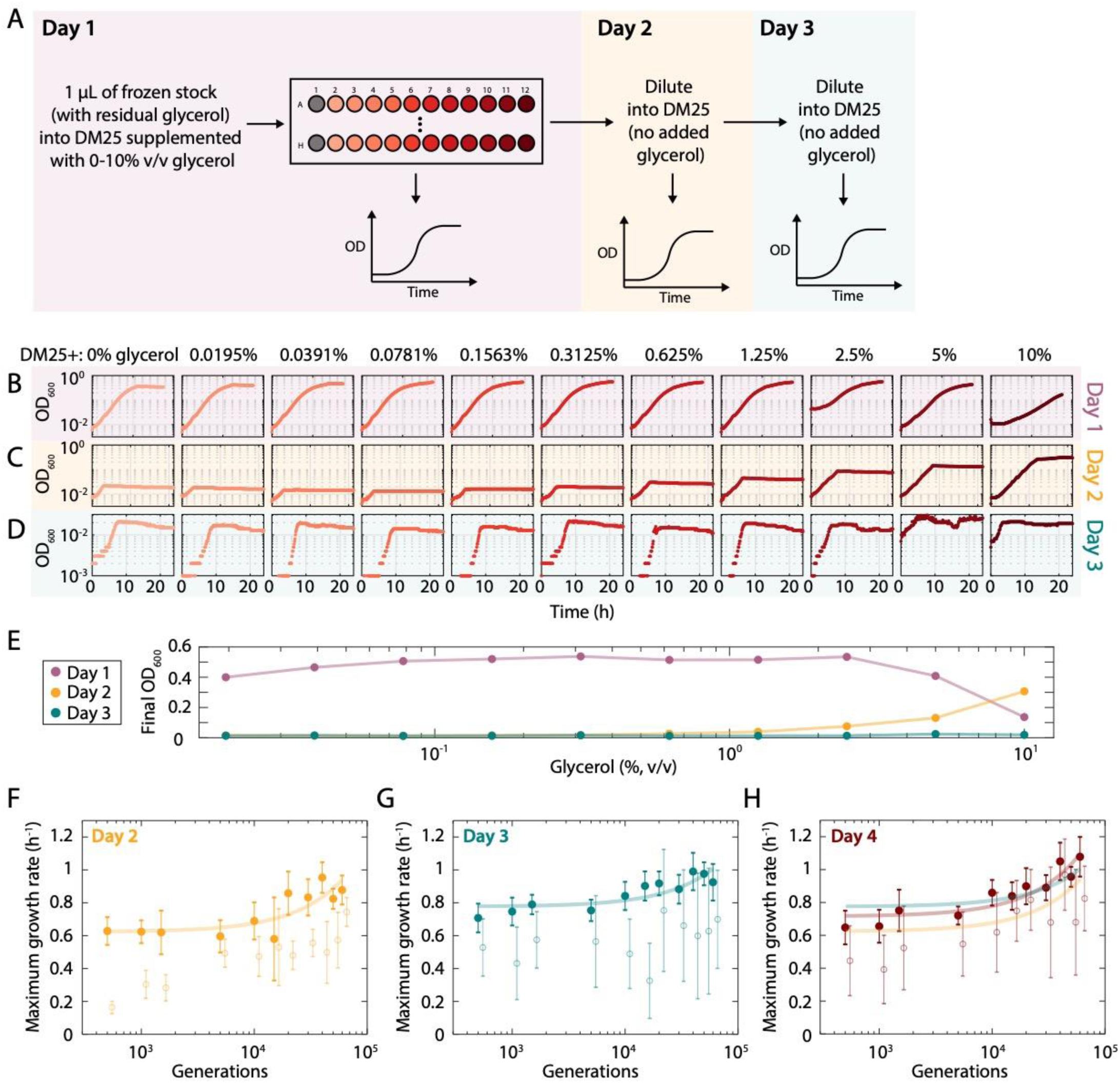
The presence of glycerol in carbon-poor medium increases the carrying capacity of that medium for *E. coli*. A) Schematic of protocol to measure the effect of glycerol on growth. *E. coli* cultures were inoculated with 1 µL of frozen stock in DM25 supplemented with various concentrations of glycerol. Growth was monitored over three passages, in which the latter two involved 1:200 dilutions into DM25 (without glycerol). B) One microliter from a thawed frozen stock (20% glycerol) of *E. coli* REL606 was grown in minimal medium (DM25, 25 mg/L glucose) with various amounts of glycerol. On day 1, the carrying capacity was much higher than would be expected for the low glucose concentration (as seen with subsequent passages). C) On day 2, the cultures were diluted into DM25, and the carrying capacity remained relatively high following the initially higher glycerol concentrations, presumably due to glycerol carryover. D) By day 3, growth curves stabilized across all concentrations, and the increase in carrying capacity due to glycerol carryover no longer occurred. Note that the *y*-axis scale is different from those in (B) and (C). E) Dependence of final OD on glycerol concentration present during growth during the first passage. Final OD was much higher on day 1 compared to day 2 and 3 for all concentrations other than 10% glycerol. On day 2, final OD increased with the glycerol concentration from the previous passage. On day 3, the dependence on glycerol concentration was gone and the final OD was very low, as expected given the carbon-poor medium. F-H) Outgrowth for multiple days minimizes glycerol carryover and enables accurate measurement of growth-rate increases at low densities in the LTEE medium. Maximum growth rates computed using well-specific blanking (filled circles) in DM25 for the Ara-1 evolved line [40] are lower and noisier during the second growth passage (E) than during the third (F) and fourth passages (G). Measurements in (F) and (G) are quantitatively similar and revealed increases in maximum growth over the course of the LTEE. The lines are linear least-squares fits to the data on day 2 (yellow), 3 (teal), and 4 (maroon). Blanking based on a randomly selected well (E-G, open circles) led to generally lower growth-rate estimates and noisier trends; these data are slightly right-shifted to avoid overlap.

Combined with our finding that growth rate can be accurately measured even at low OD values with well-specific background subtraction (Fig. 1F), we realized that we could use multiple passaging to measure growth parameters in a high-throughput manner for the LTEE strains, enabling us to determine whether they changed systematically over the course of the LTEE. We examined 12 strains sampled from the Ara-1 population through 60,000 generations [40, 41]. We diluted each culture into fresh DM25, and measured their growth curves. We then re-diluted these overnight cultures 1:200 into fresh DM25 and measured their growth curves for three more passages. All strains attained relatively high ODs in the first passage due to the residual glycerol (Fig. S2E). In the second passage, growth rates were higher for the strains from later generations, but the measurements were noisy due to variations in inoculum levels (Fig. 3E). By the third passage, the noise decreased substantially, allowing us to observe that the growth rate had gradually increased from over the 60,000 generations (Fig. 3F). By the fourth passage, maximum growth rates of the evolved strains had converged on values close to those measured the previous day (Fig. 3G), ranging from ∼0.65 h^-1^ to ∼1.08 h^-1^ from the earliest to the latest sample. Notably, the low stationary-phase density in DM25 (Fig. S2E) meant that even small amounts of noise greatly affected the measurements of growth rate; hence, well-specific blanking instead of random-well blanking was critical (Fig. 3E-G).

These data demonstrate that the serial-transfer regime of the LTEE has favored higher maximum growth rate, as predicted from theory [6] and measured previously over the first 20,000 generations [42]. That previous work, however, involved using a glucose concentration much higher than in the LTEE itself, in order to achieve OD values suitable for measuring growth rates. Our new measurements highlight the importance of accurate blanking for quantifying growth behaviors, especially at low OD values.

### *Growth in glycerol greatly increases* B. subtilis *lag time during subsequent passaging*

In addition to impacting bacterial growth in carbon-limited media, we hypothesized that glycerol could have other, species-specific effects on growth. To mimic the variable amount of frozen stock that might be used to inoculate a culture, while also avoiding confounding differences in initial cell density, we inoculated a 96-well plate with 1 µL of frozen stock of either *B. subtilis* 168 or *E. coli* MG1655 in LB supplemented with 0.1% to 10% glycerol and measured growth curves (Fig. 4A). For both species, during the first passage (day 1) all cultures exhibited approximately the same carrying capacity (Fig. 4Bi, S3Ai) and similar lag times (Fig. 4Bii, S3Aii).

**Figure 4:**
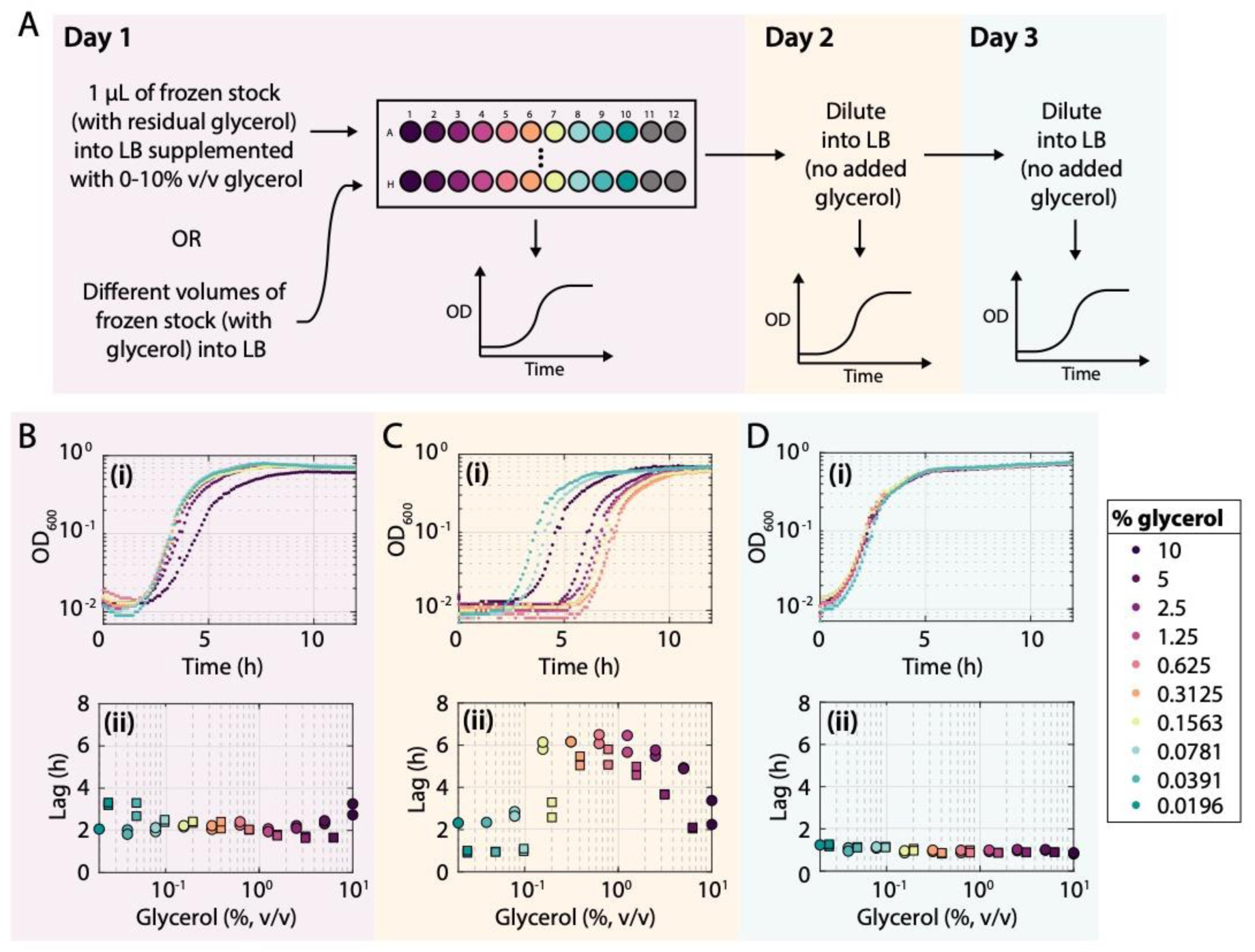
Growth with glycerol causes a large increase in lag time for *B. subtilis* in the subsequent passage. A) Schematic of protocol to measure the effect of glycerol on growth. *B. subtilis* cultures were either inoculated in LB with various amounts of a frozen stock (Fig. S3D,E) or 1 µL of frozen stock was used to inoculate LB supplemented with various concentrations of glycerol, as shown in (B-D). Growth was monitored over three passages, in which the last two followed 1:200 dilutions into LB (without glycerol). B-D) Growth curves on day 1 (Bi), day 2 (Ci), and day 3 (Di) of cultures inoculated with 1 μL of frozen stock into LB supplemented with different concentrations of glycerol (in addition to the ∼0.075% transferred with the frozen stock). During the second passage (Ci), the cultures had similar maximum growth rates and carrying capacities, but intermediate inoculation amounts led to large increases in lag time. Growth curves were essentially identical during the third passage (Di). On day 1 (Bii), lag times were roughly constant for intermediate inoculation amounts (squares) or glycerol concentrations (circles), but lag times increased dramatically on day 2 (Cii); by day 3 (Dii) there was little difference in lag times across glycerol concentrations. These data indicate that glycerol causes the long-lag phenotype. Similar results were obtained when inoculating with different volumes of frozen stock (Fig. S3).

At high levels glycerol causes catabolite repression and supports growth rates similar to glucose [43]. Hence, we speculated that glycerol could alter the physiological state of cells as they progress through another passage. To determine whether subsequent growth was affected by the prior presence of glycerol, we diluted all cultures 1:200 into fresh LB (without adding glycerol) and monitored growth in a plate reader for a second passage (day 2) (Fig. 4Ci, S3Bi). For intermediate concentrations between 0.16% and 5%, the lag time of *B. subtilis* cultures during this second passage increased to 6 h (Fig. 4Cii). This increased lag was specific to *B. subtilis* (Fig. S3Bii, S3Eii, S3Hii), and it went away during a third passage (Fig. 4D). Similar behavior was seen when inoculating a 96-well plate with fresh LB with various volumes (0.1-100 µL) of thawed frozen stocks of *B. subtilis* 168 (Fig. 4A, S3D-F). These data indicate that the second passage after revival of *B. subtilis* from a frozen stock is very sensitive to the glycerol concentration during initial inoculation.

To confirm that cellular responses to freezing were not required for the increase in lag time, we used 1 µL of an overnight culture grown from frozen stock to inoculate LB supplemented with 0.1% to 10% glycerol; these cultures were now removed from freezing by a 24-h passage (Fig. S4A, purple). During the passage after glycerol addition, cells exhibited the expected increases in lag time (Fig. S4A, purple). Furthermore, when we inoculated the initial culture from a colony (instead of a frozen stock) into LB supplemented with glycerol, we saw similar increases in lag time on the subsequent passage (Fig. S4A, yellow) as from inoculating in varying amounts of glycerol (Fig. 4C; S4A, teal) or different amounts of frozen stock (Fig. S3E). These findings suggest that regrowth from frozen stock displays little to no sign of cell death relative to regrowth from a colony. The long-lag phenotype in the presence of glycerol for *B. subtilis* is distinct from the effects we observed in *E. coli* (Fig. 3B-E), in which the glycerol was metabolized and thus led to higher carrying capacities. Altogether, these data show that growth of *B. subtilis* 168 in glycerol can cause dramatic lag time increases during the subsequent passage for intermediate concentrations of glycerol. They highlight the importance of controlling for glycerol levels during high-throughput growth assays, which can be readily achieved by performing an additional passage.

### Glycerol has dramatic and varied effects on B. subtilis single-cell growth

We were surprised by the increased lag times in *B. subtilis* growth after passaging with intermediate glycerol concentrations, and sought to investigate the cell physiology underlying this phenomenon. We used time-lapse microscopy to monitor the growth of cells on LB agarose pads following growth in liquid LB at the glycerol concentrations associated with the longest population lag times (Fig. 4C). For cells grown in liquid LB with 0.3125% glycerol (∼6 h lag time, Fig. 4Cii), we imaged an entire ∼2 mm-diameter spot by capturing a grid of 144 overlapping fields of view. Of the ∼10^4^ cells in one spot, only a single cell grew, and its descendants took over the entire spot over 12 h of imaging (Fig. 5B). That cell exhibited growth as soon as imaging began, and after 1.7 h its lineage exhibited a doubling time of ∼20 min (Fig. 5C), indicating the extreme heterogeneity in this population. In other spots (*n* = 2), we observed no growth of any cells. Such extreme bottlenecks should be avoided for most applications.

**Figure 5:**
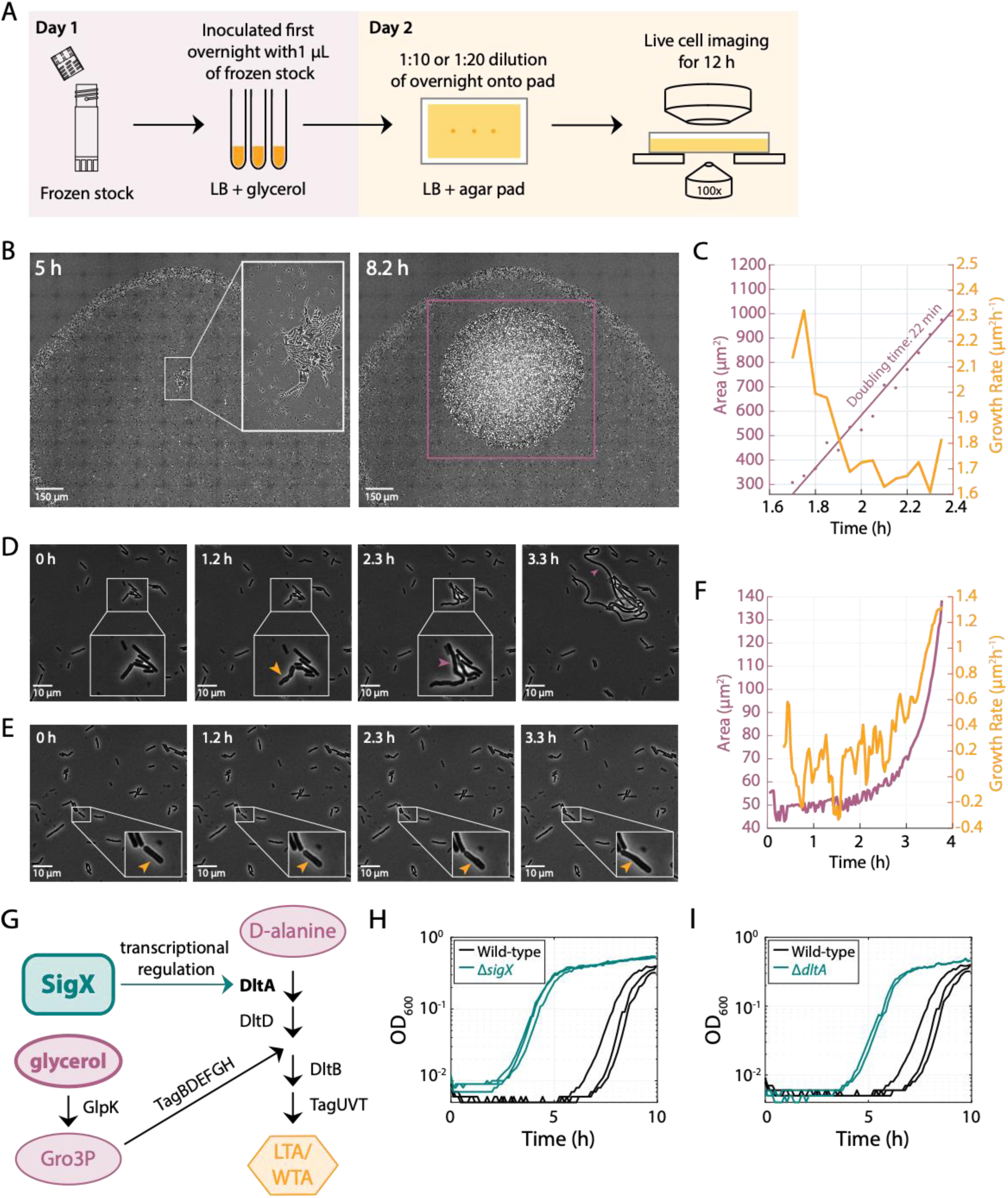
Growth of *B. subtilis* in intermediate glycerol concentrations results in highly heterogenous single-cell phenotypes during the subsequent passage. A) Schematic of protocol for imaging single-cell growth during the passage after growth in glycerol. B) Time-lapse images of a stitched set of 144 fields of view covering the entire spot of cells on an LB agarose pad after a passage in LB + 0.3125% glycerol. Inset, enlarged view of the only cell that exhibited growth across the entire spot. The purple box at right highlights the resulting microcolony at 8.2 h. C) Quantification of cell area (purple) and instantaneous growth rate (yellow) of the initially sole growing cell in (B) finds a maximum growth rate corresponding to a doubling time of 22 min. D) Time-lapse of cell growth on an LB agarose pad after growth in liquid LB + 0.625% glycerol. The cell (inset, enlarged) started growing immediately, and bulged (yellow arrow) and filamented for ∼3 h (purple arrow). E) Time-lapse of cell growth (yellow arrow; inset, enlarged) on an LB agarose pad after growth in liquid LB + 0.625% glycerol; this cell did not start growing until after ∼2 h. F) Quantification of cell area (purple) and instantaneous growth rate (yellow) of the cell in (E), which had a longer lag and slower growth rate than the cell in (C). G) Schematic of regulatory pathway for biosynthesis of LTA and WTA involving SigX and DltA. SigX transcriptionally regulates *dltA* and 47 other genes, some of which are involved in biosynthesis of teichoic acids. Glycerol is converted to *sn*-Glycerol 3-phosphate (Gro3P); Gro3P, in conjunction with an intermediate product made by DltA, is necessary for the production of teichoic acids. H,I) Deletions (teal) of *sigX* (H) and *dltA* (I) resulted in a shorter lag than wildtype (black).

For cells grown in LB with 0.625% glycerol (∼6 h lag time, Fig. 4Cii), we again observed a small fraction of growing cells (<3%) (Fig. 5D). The cells that grew showed multiple phenotypes. Some cells started growing immediately (Movie S1); some initially bulged along the cell body and then filamented for ∼2.5 h before the first division (Fig. 5D). Other cells did not grow for >2 h (Fig. 5E), and thereafter grew more slowly than normal (Fig. 5F, Movie S2). A third subset periodically shrank and had a high death rate (Movie S3). For one cell, no growth was observed for the first 3 h; afterwards it engaged in short phases of growth and shrinking, with small blebs forming during growth (Fig. S5A). It eventually divided, and many of its progeny also exhibited periodic shrinking (Movie S3) and death through explosive lysis (Fig. S5A). Thus, growth in glycerol clearly disrupts cell shape as well as growth out of stationary phase in multiple ways.

### A screen links glycerol-induced lag to genes involved in teichoic acid synthesis

The discovery of *B. subtilis*’s long lag and consequent fitness defect induced by intermediate glycerol concentrations (Fig. 4) presented the opportunity to identify genetic determinants of this phenotype, as mutants with a shorter lag would be enriched in the population. To gain insight into the underlying mechanism, we carried out an unbiased genetic screen by creating six independent pooled libraries of transposon mutants (Methods) [44] in the wild-type strain. We grew the libraries in LB + 1.25% glycerol, a concentration that induced a long lag time in wildtype (Fig. 5Eii). Further passaging of the libraries once in LB + 1.25% glycerol and once more in LB revealed two libraries that evolved shorter lag times (Fig. S5B). We isolated single colonies and verified that they had a similar phenotype to the library from which they were isolated (Fig. S5C). Sequencing their transposon insertion sites identified two mutations: in *sigX*, which encodes a sigma factor that regulates modification of the cell envelope and resistance to cationic antimicrobial peptides [45], and in the start codon of *dltA*, which encodes a D-alanine ligase required for modification of wall teichoic acids (WTA) and lipoteichoic acids (LTA) [46]. Note that *dltA* is part of the *sigX* regulon (Fig. 5G) [45]. We verified these hits by reintroducing each transposon insertion into the parental strain (Fig. S5D,E) and by deleting the gene (*sigX* or *dltA*), and then showing that these constructs exhibited the same reduction in lag (Fig. 5H,I). The thick peptidoglycan cell wall of Gram-positive bacteria is intercalated with wall teichoic acids, which are covalently bound to peptidoglycan, and lipoteichoic acids, which are tethered to the membrane by a lipid anchor [47]. Production of both wall teichoic acids and lipoteichoic acids requires substantial amounts of glycerol [48]. Thus, it appears that teichoic-acid production is linked to the glycerol-dependent long-lag phenotype in *B. subtilis*.

## Discussion

As microbiology research has expanded and flourished, so has the appreciation of the sensitivity of microbial physiology and cellular structure to environmental conditions. Uncovering the details of these sensitivities will be critical to quantitative understanding of growth behaviors across microbial strains and species, as will establishing the resolution and robustness of the equipment used to measure growth. We have described a general strategy for measuring an instrument’s limit of detection and range of linearity, and demonstrated that, with proper protocols, a wide range of growth behaviors can be accurately quantified.

The dependence of maximum growth rate on initial cell density presents complications similar to the antibiotic inoculum effect, whereby the sensitivity to certain drugs increases at lower cell density [49]. Comparisons of growth rate and lag time between strains would ideally employ similar initial cell concentrations. However, fulfilling such a condition can be challenging due to strain differences in cell shape [50], yield in a given medium, and cell survival in and recovery from stationary phase. Moreover, some species may have growth dynamics and carrying capacities that are inoculum-dependent. Given these complications, the acquisition of growth curves across a range of initial densities to map the range of growth behaviors would enhance the ability to compare strains and species. Our model based on ordinary differential equations is general and hence can be used for many microbes; in particular, it can help to correct for differences in growth rate due to variation in initial inoculum size. It is also important to note that waste accumulation should be mathematically equivalent to nutrient depletion if waste products inhibit the uptake of some nutrients, which means our model is even more broadly applicable. However, other factors may prove important for modeling growth curves, such as pH changes that are known to inhibit growth [51].

Although spectrophotometers that read microtiter plates are quite sensitive to small changes in OD (Fig. 1D,E), our analyses establish that it is critical to minimize noise that introduces complexities when calculating growth metrics; this noise minimization can be best achieved by blanking each individual well separately. This modification to protocols was relatively straightforward, and it positioned us to quantify the contribution of increased growth rate to the fitness gains observed in the LTEE with *E. coli* (Fig. 3). In particular, this method allowed us to show unequivocally that faster exponential growth had been selected even at the low glucose concentration and consequently low OD values of the LTEE. The ∼66% increase in maximum growth rate that we measured is reasonably close to the ∼70% increase in relative fitness obtained through competing late-generation samples against their ancestor [39]; that relative fitness is calculated as the ratio of the realized growth rates of the evolved and ancestral bacteria over a full 24-hour transfer cycle including lag, growth, and stationary phases. Thus, other growth parameters can also affect relative fitness including differences in lag time and carrying capacity [52], which complicates a direct comparison between maximal growth rate and relative fitness. In addition, crossfeeding interactions have evolved in some LTEE populations, and these interactions may affect the post-maximum growth rates of the competitors as cells exhaust the limiting glucose, consume byproducts, and transition into stationary phase [53-55]. In any case, our new protocol and growth-rate measurements demonstrate the value of subtracting the blank of each well to minimize noise, especially at low OD values. These growth-rate data also imply that utilization of glucose has become much faster over the LTEE, and more generally it may be possible to evolve many bacterial species to grow at higher rates in specific nutrient conditions. Given the correlation between cell size and fitness in the LTEE [56], future studies might use these strains to explore whether the “Growth Law” that links nutrient-dependent growth rate and cell size [57] has changed over the course of evolution. In fact, it was previously shown that that the relation is not constant between the ancestor and an evolved strain isolated after just 2,000 generations [58].

The dramatic increases in lag time that we observed in *B. subtilis* (Fig. 4C) indicate that glycerol can have lasting physiological effects that impede the future growth of cells. We did not observe this lag phenotype in *E. coli* (Fig. S3B). This difference is consistent with the requirement of Gram-positive bacteria for glycerol to synthesize teichoic acids, which they incorporate into the cell envelope. Without multiple dilutions to mitigate the glycerol-induced long-lag phenotype, a severe bottleneck in which very few cells are responsible for outgrowth can occur (Fig. 5B), which may complicate interpretation. The conventional approach to streaking single colonies before starting liquid cultures avoids this problem (Fig. S4A,B), likely due to the extreme dilution of the glycerol concentration. While streaking may be prohibitive for large strain collections or certain species and communities, washing the initial inoculum to remove the glycerol is also sufficient (Fig. S2C,D).

Our transposon mutagenesis screen linked the glycerol-induced long-lag phenotype in *B. subtilis* to the incorporation of glycerol into lipoteichoic acids (Fig. 5H,I). Our data also revealed several unusual physiological and morphological impacts of glycerol in the subsequent passage including filamentation (Fig. 5D), bulging (Fig. 5E), lysis (Fig. S5A), and repeated cycles of growth and shrinkage (Fig. S5A). Some of these phenotypes are consistent with previous observations connecting lipoteichoic-acid synthesis with cell division [59]. Moreover, the arrest of growth in the vast majority of cells (Fig. 5E) indicates that their physiological state was sensitized by previous exposure to even low amounts of glycerol, such that growth remained inhibited even after glycerol was no longer present. This phenomenon connects teichoic-acid synthesis to growth inhibition and lag phase for the first time, and it points to a severe bottleneck that might cause additional experimental complications. This surprising phenotype also highlights the potential for other physiological history-dependent effects on growth.

In the process of dissecting the seemingly simple process of measuring bacterial growth, we developed a refined protocol that enabled precise, quantitative measurements. This precision, in turn, led to the discovery of new biological phenomena. The microbial world is stunningly diverse, and we currently know very little about the growth kinetics of the vast majority of microbes. Our work provides a powerful framework to analyze the growth characteristics of microbial species in high-throughput assays.

## Methods and Materials

### Strains and media

Table S1 lists the strains and their genotypes used in this study. Strains were grown in LB (Lysogeny broth with 10 g/L tryptone, 5 g/L NaCl, and 5 g/L yeast extract) or DM25 (Davis minimal broth [60] supplemented with 2 × 10^−3^ g/L thiamine hydrochloride and 25 mg/L glucose [37]). Both media were supplemented with 0-10 % (v/v) glycerol when indicated. When stated, antibiotics were used as follows, unless indicated otherwise: kanamycin (5 *μ*g/mL) and MLS (a combination of 0.5 *μ*g/mL erythromycin and 12.5 *μ*g/mL lincomycin).

### Bacterial growth

For all glycerol experiments, to avoid potential issues associated with accumulated damage over time spent in the freezer, we first made new frozen stocks of *E. coli* MG1655 and *B. subtilis* 168 by growing a single colony overnight in LB and then rapidly freezing a 1:1 mixture of the culture with 50% glycerol in a −80 °C freezer. All strains (Table S1) were inoculated from −80 °C freezer stocks into 200 µL of the medium of interest in shallow 96-well plates (Greiner) and grown overnight for 18 h while shaking at 37° C. Overnight cultures were diluted 1:200 into fresh medium for plate-reader experiments, and either 1:10 or 1:20 into fresh medium for microscopy.

### Plate-reader absorbance measurements

A shallow 96-well plate was filled with fresh medium, and a plate seal (Excel Scientific) with laser-cut holes was placed on top. The background absorbance at 600 nm (OD_600_) was measured for 30 min to allow the readings to stabilize. Overnight cultures were diluted 1:200 into this plate using the same plate seal and grown with shaking at 37 °C in a BioTek Epoch 2 plate reader for 18-24 h. OD_600_ was measured at 7.5-min intervals. For the resulting growth curves, the slope over a sliding window (for smoothing) was computed to determine the instantaneous growth rate, and from that, the lag time (defined as the time to half-maximal growth rate) was calculated. Additionally, the natural logarithm of OD_600_ was fit to the Gompertz equation [12] to quantify lag time and maximal growth rate. The two methods for calculating lag times and maximum growth rates were highly correlated (Fig. S6) for growth curves of *E. coli*. The first method was more appropriate for growth curves that did not reach saturation or exhibited a distinct shape (e.g., due to a diauxic shift). Thus, we used the first method to quantify lag time and maximal growth rate, except as noted.

### Microscopy

Cells were diluted 1:10 or 1:20 (depending on the overnight culture OD) into fresh LB. For time-lapse imaging, 1 µL of cell culture was placed onto a large pad (the size of a 96-well plate) composed of 1.5% agar + LB. Such a large pad was used to avoid oxygen depletion in the pad over the course of imaging, because *B. subtilis* cells lyse in oxygen-limited conditions [8]. The cells were imaged using a Nikon Eclipse Ti-E inverted fluorescence microscope with a 100X (NA 1.40) oil-immersion objective, and integrated using µManager v. 1.3 [61]. Cells were maintained at 37 °C during imaging with an active-control environmental chamber (HaisonTech).

### Strain construction

We constructed strains to study the role of LTA synthesis in glycerol-induced lag using SPP1 phage transduction [62]. The donor strain was grown for >6 h in TY medium (LB supplemented with 0.01 M MgSO_4_ and 0.1 mM MnSO_4_ after autoclaving). Ten-fold dilutions of SPP1 phage were added to the culture, and 3 mL of TY soft (0.5%) agar were then mixed with the cell/phage mixture and poured over a TY plate (1.5% agar) before overnight incubation. We chose a plate that exhibited nearly complete clearing of cells, thus without many phage-resistant mutants. Five milliliters of TY medium were added to this plate and a 1-mL filter tip was used to scrape up the soft agar. This soft agar/liquid mix was filtered through a 0.4-µm filter (Fisher Scientific). The phage were added to a stationary-phase culture (grown for 6-10 h) of the recipient strain (10 μL undiluted phage + 100 μL recipient cells, and optionally 900 μL TY medium) in TY medium and incubated at 30 °C for 30 min, then plated onto LB + antibiotic and 0.01 M sodium citrate (sodium citrate was omitted for MLS selection). Plates were incubated for 24 h at 30 °C and transductants were streaked for single colonies to eliminate the phage.

The back-crossed transposon mutagenesis strains and deletion strains (Δ*sigX* and Δ*dltA*) were constructed via double-crossover integration by transformation of genomic DNA using standard transformation procedures [63, 64]. We performed genomic DNA extractions of the original transposon mutagenesis strain, BKK23100 (*sigX*), and BKK38500 (*dltA*) using the Promega Wizard kit.

### Mariner transposon library construction

The mariner transposon was transduced into the parent strain as described above. The mariner plasmid has a temperature-sensitive origin of replication, a transposase, and a transposon that is marked with kanamycin resistance; the backbone plasmid is marked with MLS resistance [65]. Before transduction of the mariner transposon, the parent strain (CAG74168) was streaked for single colonies on MLS plates at 30 °C. One colony for each library was grown overnight in 3 mL LB + kanamycin in a roller drum at room temperature. Transposon-insertion libraries were created by plating 10-fold dilutions of the cultures on LB + kanamycin plates prewarmed to 37 °C and incubating them overnight at 37 °C. The dilution that contained very dense colonies was scraped to create the library, and the higher dilutions were used to estimate the number of transposon mutants per library.

### Transposon library screen

The transposon library was screened by using the entire library to inoculate a 5 mL LB + 1.25% glycerol culture. This culture was diluted 1:200 after 18 h of growth into 200 µL of LB + 1.25% glycerol, grown for 18 h with shaking at 37 °C, and diluted 1:200 into LB. After two passages of growth in LB + glycerol and one passage in LB, the libraries with a shorter lag time were streaked for single colonies on LB plates. Five single colonies were picked for each positive hit and used to inoculate a 200-µL LB culture. After 18 h, the culture was diluted 1:200 into LB + 1.25% glycerol. The culture was grown for 18 h and diluted 1:200 into LB. After three passages, the positive hits were grown in 5 mL of LB for genomic DNA extraction.

### Mapping transposon insertions using inverse PCR

Genomic DNA was extracted using the Promega Wizard Genomic DNA purification kit, digested with Sau3AI at 37 °C for 90 min, and then heat-inactivated at 65 °C for 20 min. The reaction mixture contained 15.5 μL milliQ H_2_O, 2 μL NEBuffer1.1, 2 μL digested genomic DNA, 2 μL 10X bovine serum albumin, and 0.5 μL Sau3AI. The digested DNA was ligated using T4 ligase at room temperature for at least 1 h; the reaction mixture contained 2 μL T4 DNA Ligase Buffer, 15.5 μL milliQ H_2_O, 2 μL digested DNA, and 0.5 μL T4 DNA ligase. Inverse PCR was carried out with the ligated DNA using Phusion polymerase with the primers IPCR1 (5’-GCTTGTAAATTCTATCATAATTG-3’) and IPCR2 (5’-AGGGAATCATTTGAAGGTTGG-3’). Each reaction contained 33 μL milliQ H_2_O, 10 μL 5X HF buffer, 2 μL ligated DNA, 2 μL IPRC1, 2 μL IPCR2, 1 μL 10 mM dNTPs, and 0.2 μL Phusion Polymerase. The PCR program was as follows: 98 °C for 30 s, 30 cycles of [98 °C for 10 s, 58 °C for 30 s, 72 °C for 60 s], 72 °C for 10 min, and hold at 4 °C. PCR products were gel-purified and sequenced using the IPCR2 primer. The sequences were mapped onto the *B. subtilis* 168 genome using BLASTN.

### Data availability

All data used in this manuscript are growth curves, time-lapse microscopy images, or transposon sequencing. All data are available upon request from the corresponding author.

## Acknowledgments

The authors thank the Huang lab for useful discussions. This work was supported by a Stanford Graduate Fellowship (to E.A.), an NSF Graduate Research Fellowship (to S.C.), an NIH Ruth Kirschstein Award F32-AI133917 (to M.R.), a Bio-X Graduate Fellowship (to A.A.-D.), an HHMI International Student Fellowship (to A.A.-D.), an Agilent Graduate Fellowship (to H.S.), a Stanford Interdisciplinary Graduate Fellowship (to H.S.), a James McDonnell Postdoctoral Fellowship (to H.S.), NSF grant DEB-1951307 (to R.E.L.), USDA Hatch grant MICL02253 (to R.E.L.), NSF CAREER Award MCB-1149328 (to K.C.H.), and the Allen Discovery Center at Stanford on Systems Modeling of Infection (to K.C.H.). K.C.H. is a Chan Zuckerberg Biohub Investigator.

## Supplementary Figure Legends

**Supplementary Figure 1:**
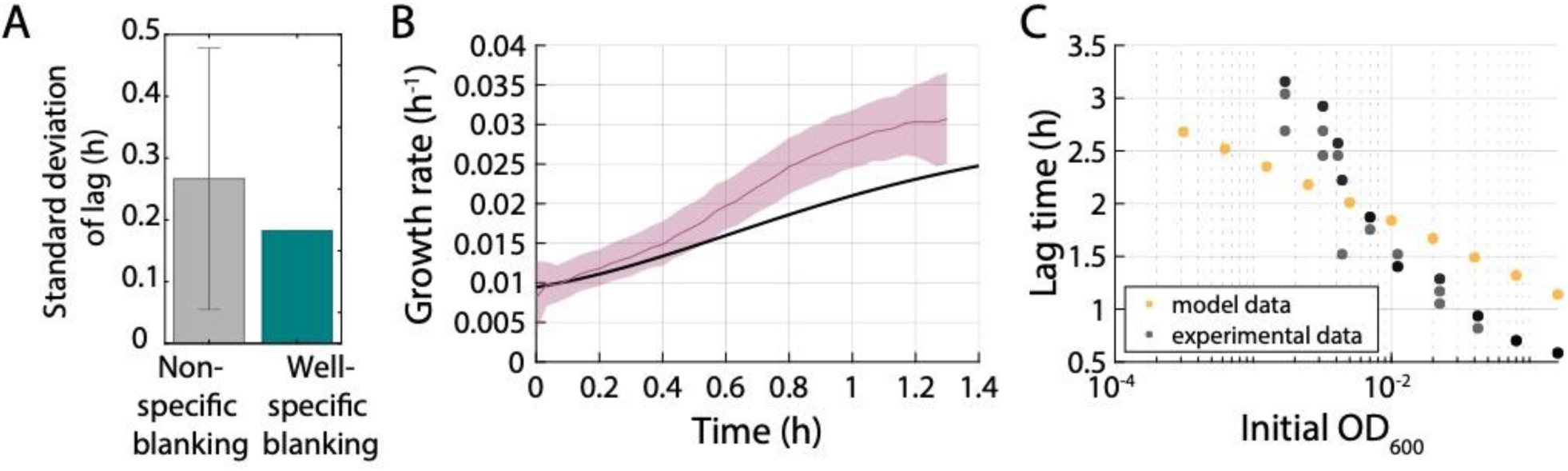
Fitting parameters for a model of bacterial population growth that depends on inoculum size, and comparison of this model to experimental data. A) Well-specific blank subtraction (teal) decreases the standard deviation in the estimate of lag time compared to subtracting a randomly selected blank in a 96-well plate (gray). B) The instantaneous growth rate of *E. coli* MG1655 cells extracted from single-cell imaging on LB + 1% agarose pads during emergence from stationary phase increases over the first hour. Shaded region indicates ±1 standard deviation. The black line is the Gompertz model with *λ* = 0.8 h^−1^, *δ* = 0.5 h. C) Simulated growth curves based on Eqs. 4-6 in the main text capture, at least roughly, the decrease in lag time with increasing initial inoculum density.

**Supplementary Figure 2:**
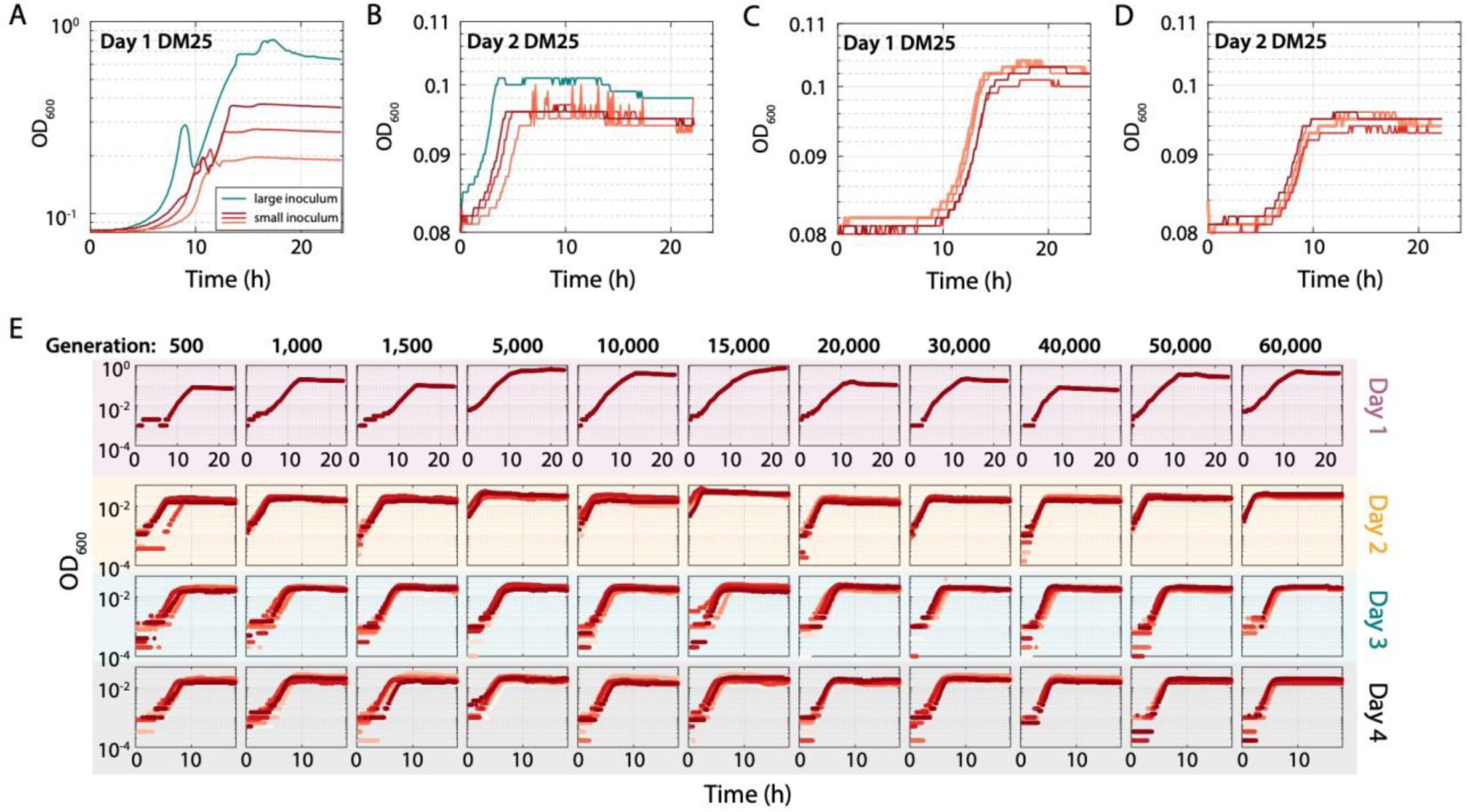
Volume from a frozen stock used to inoculate a culture affects carrying capacity in low-resource medium. A) Various volumes of a frozen stock of *E. coli* REL606 were used to inoculate cultures grown in DM25. Dark to light red indicate three replicates of a small inoculum size (small but inexact volumes were picked from the frozen stock), while teal indicates an intentionally much larger inoculum volume. The large inoculum (teal) led to a higher final OD on the first day of growth. B) When the cultures in (A) were diluted 1:200 into fresh DM25 and regrown for 24 h, the higher final ODs on day 1 translated into slightly higher ODs (teal) even on day 2. C) *E. coli* REL606 cultures were inoculated by resuspending a small volume of the frozen stock in 200 μL of DM25, spinning down for 5 min at 8000*g*, removing the supernatant, and resuspending the pellet in 200 μL of DM25 for growth. The curves represent four technical replicates. D) When the cultures in (C) were diluted 1:200 into fresh DM25 and regrown for 24 h, the replicates behaved similarly and had a slightly lower final OD than the large inoculum in (B) (teal). E) The *E. coli* Ara-1 line evolved a higher maximum growth rate over the course of 60,000 generations. One microliter of frozen stock (20% glycerol) of each clone (shown by generation sampled) from the *E. coli* Ara-1 line [40] was grown in DM25 with various amounts of glycerol across four 24-h passages. On day 1, the carrying capacity was much higher than was achieved in later days in the low-glucose DM25 medium. The populations were diluted 1:200 on subsequent days into DM25 and growth curves were monitored in a plate reader. The maximum growth rate extracted from the curves is more accurately measured on Days 3 and 4, and it increases over evolutionary time (Fig. 4). All curves are shown after subtracting the blank value of the corresponding well.

**Supplementary Figure 3:**
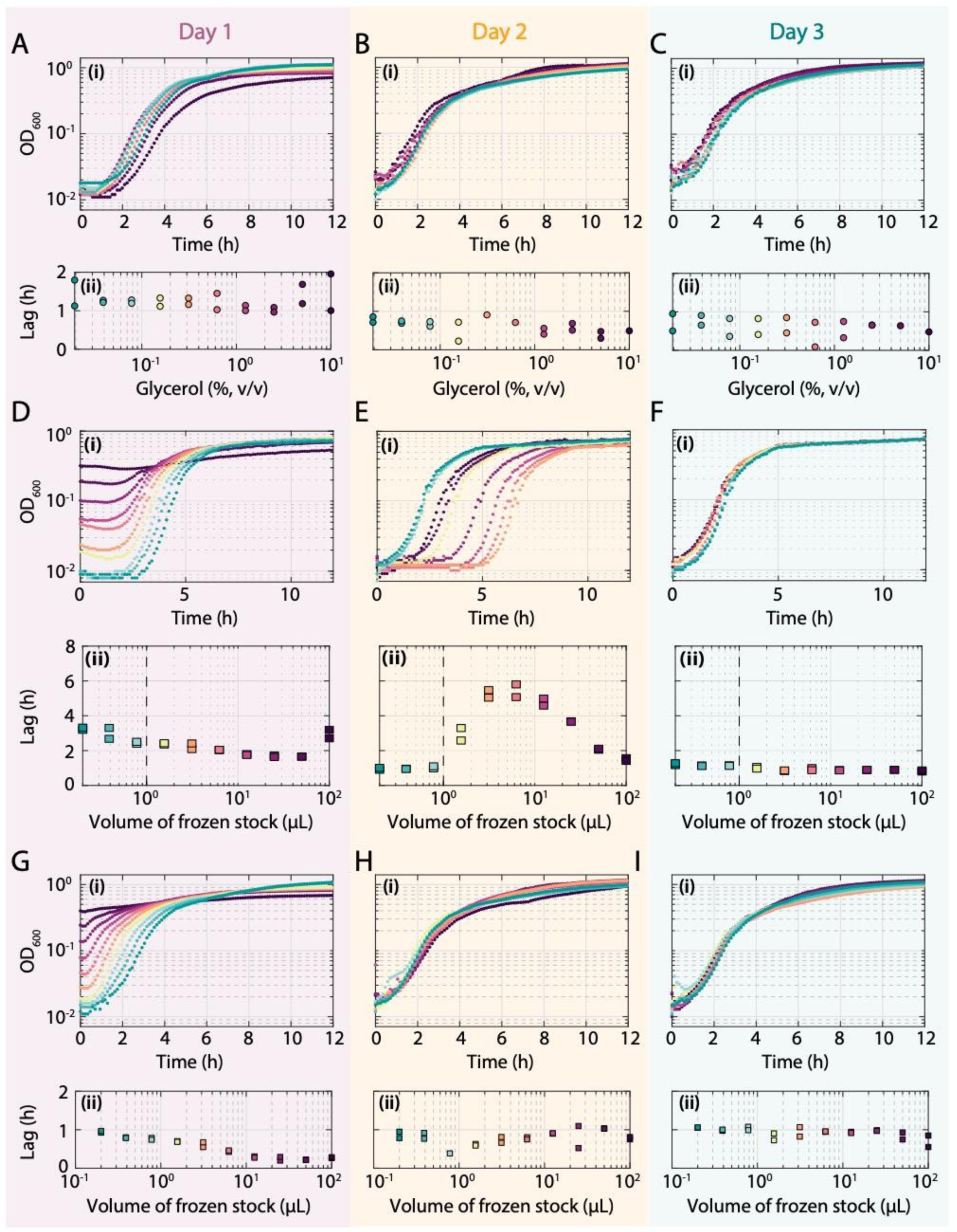
Glycerol has little effect on *E. coli* growth in nutrient-rich LB. A-C) Growth curves of *E. coli* MG1655 in LB supplemented with varying amounts of glycerol on day 1 (Ai), diluted 1:200 into LB on day 2 (Bi), and diluted again 1:200 into LB on day 3 (Ci). There were slight glycerol-dependent effects on day 1 (Ai), but growth curves on days 2 (Bi) and 3 (Ci) were nearly identical for all cultures. Estimates of lag time from the growth curves are shown in (Aii), (Bii), and (Cii). Lag times on day 1 (Aii) were longer than on days 2 (Bii) and 3 (Cii), which were quantitatively similar across glycerol concentrations. D-F) Growth curves on day 1 (Di), day 2 (Ei), and day 3 (Fi) of cultures inoculated with various volumes of frozen stock. During the second passage (Ei), the cultures had similar maximum growth rates and carrying capacities, but intermediate inoculation amounts led to much longer lag times Growth curves were essentially identical during the third passage (Fi). Estimates of lag time from the growth curves are shown in (Dii), (Eii), and (Fii). On day 1 (Dii), lag times were roughly constant for intermediate inoculation amounts of glycerol, but they increased dramatically on day 2 (Eii), before becoming uniform and independent of glycerol concentration on day 3 (Fii). These results are consistent with the increase in lag shown in Fig. 4C. G-I) Growth curves of *E. coli* MG1655 in LB supplemented with various volumes of frozen stock on day 1 (Gi), diluted 1:200 into LB on day 2 (Hi), and diluted again 1:200 into LB on day 3 (Ii). Inoculum volume affected initial OD, maximum growth rate, and lag time on day 1 (Gi), but growth curves on day 2 (Hi) and day 3 (Ii) were nearly identical for all cultures. Estimates of lag time from the growth curves are shown in (Gii), (Hii), and (Iii). Lag times on days 2 (Hii) and 3 (Iii) were quantitatively similar across glycerol concentrations.

**Supplementary Figure 4:**
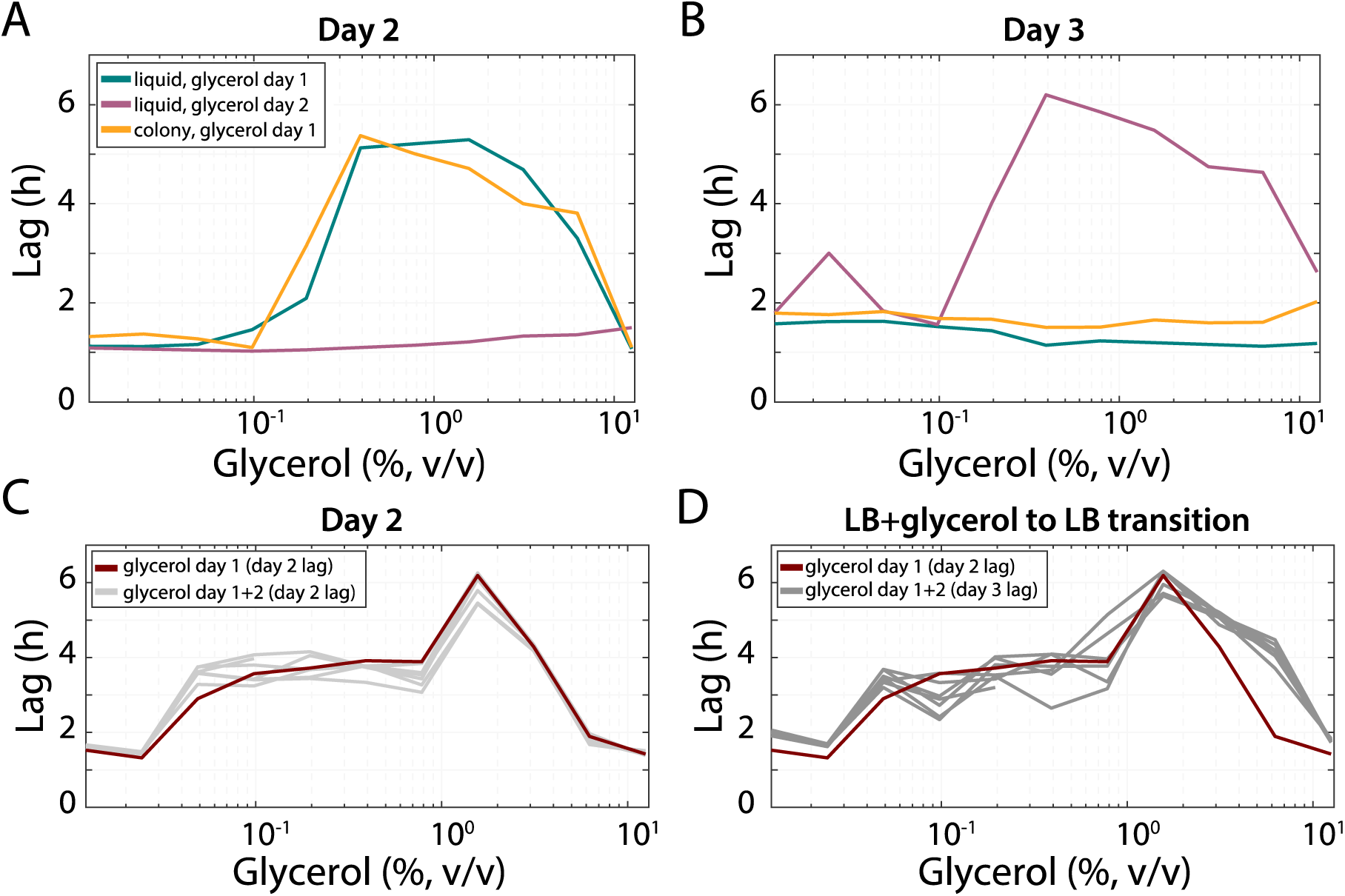
Glycerol causes the long-lag phenotype in *B. subtilis*. A) *B. subtilis* 168 was inoculated from a liquid culture (teal) or a colony (yellow) and grown in LB supplemented with various concentrations of glycerol. In both cases, there were similar lag increases on day 2 after 1:200 dilution and growth in LB. By contrast, there was no long-lag phenotype when cultures (initially inoculated with 1 µL of frozen stock on day 1) were grown with glycerol on day 2 rather than day 1 (purple). B) During the next dilution into LB (day 3), the cultures in (A) inoculated with liquid (teal) or colonies (yellow) exhibited short lags, while the glycerol cultures on day 2 (purple) displayed long lags, as expected due to growth in glycerol on the previous day. C,D) Growth of *B. subtilis* 168 in LB supplemented with glycerol for two consecutive days caused long lags at intermediate glycerol concentrations on day 2 with glycerol (C, gray) and day 3 without glycerol (D, gray), similar to one day of growth in glycerol (maroon).

**Supplementary Figure 5:**
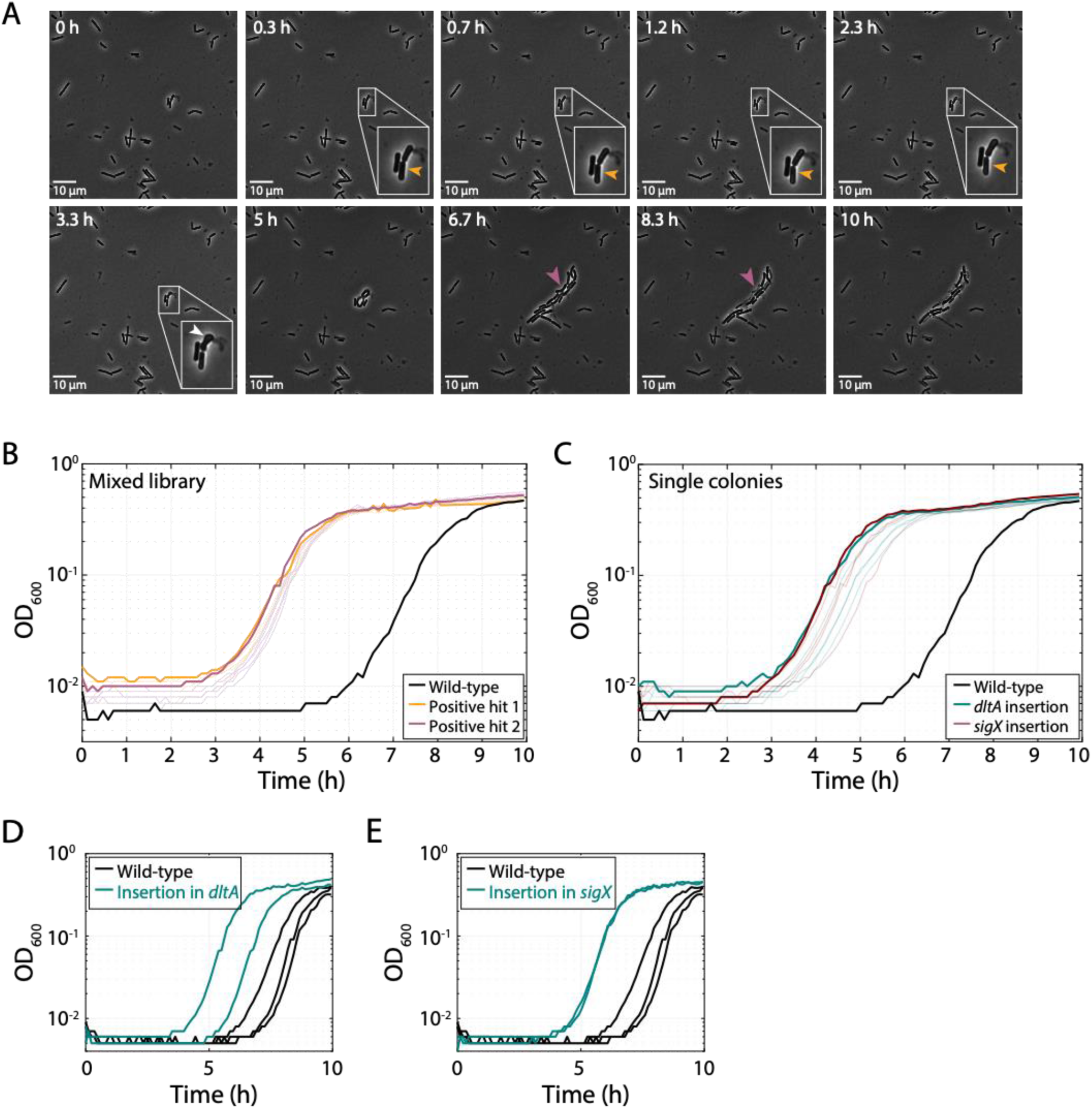
Mutations in genes involved in teichoic-acid production shorten glycerol-induced lag times in *B. subtilis*. A) Time-lapse images of a cell on an LB agarose pad after growth in liquid LB + 0.625% glycerol; this cell did not grow for >3 h, then engaged in short bursts of growth and shrinking (yellow arrows). Small blebs formed on another cell during growth (white arrows), and many of its progeny also exhibited periodic shrinking and died by explosive lysis (purple arrows). B) A transposon mutagenesis screen yielded two positive hits (yellow and purple) of libraries with shorter lag times during passaging after growth in LB + 1.25% glycerol than the parental, wild-type strain (black). Light purple and light yellow curves show 6 biological replicates. C) Single colonies isolated from the two libraries in (A) had shorter lag times during passaging after growth in LB + 1.25% glycerol than the parental, wild-type strain (black). Sequencing revealed that the colonies have insertions in *dltA* (teal) and *sigX* (maroon), both of which are linked to lipoteichoic-acid synthesis. Light teal and light maroon curves show 5 biological replicates. D,E) Reintroduction of the transposon insertions (teal) in *dltA* (D) and *sigX* (E) from the positive hits of the screen in (C) resulted in a shorter lag than wildtype (black). Two independent colonies were tested for each insertion.

**Supplementary Figure 6:**
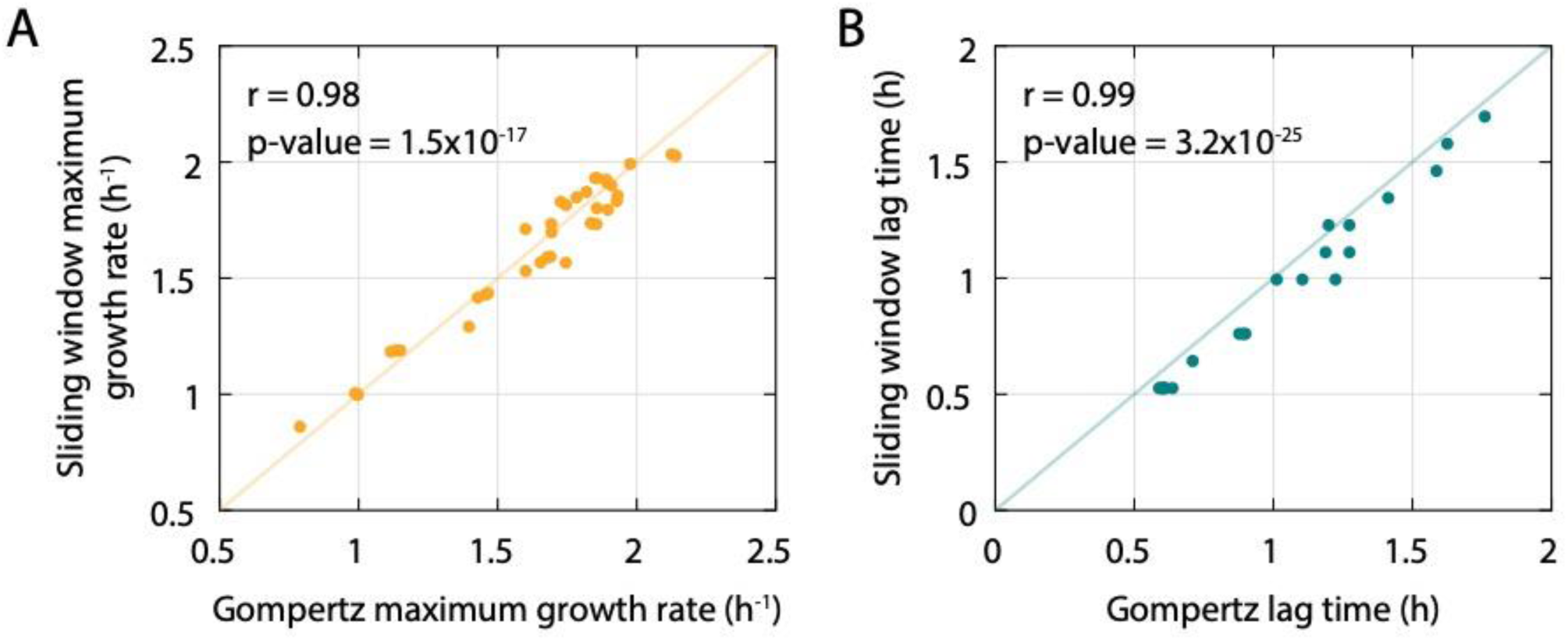
Fitting a Gompertz function and a linear model to a sliding window of log-transformed OD values provides nearly identical estimates of growth parameters in *E. coli*. A) Maximum growth rates calculated using a Gompertz fit or using a sliding window were highly correlated (*x* = *y* line in yellow). B) Lag time calculated using a Gompertz fit was highly correlated (*x*=*y* line in teal) with the time to reach half-maximum growth rate calculated using a sliding window fit.

## Movie Legends

**Movie S1: A small fraction of cells exhibit bulging and filamentous growth after overnight passage in LB + 0.625% glycerol, related to Fig. 5D.** Wild-type *B. subtilis* 168 cells from an 18-h culture in LB + 0.625% glycerol were spotted onto an LB-agarose pad and imaged every 2 min at 37 °C. This experiment was done in tandem with Movies S2 and S3.

**Movie S2: A small fraction of cells exhibit long lags after overnight passage in LB + 0.625% glycerol, related to Fig. 5E.** Wild-type *B. subtilis* 168 cells from an 18-h culture in LB + 0.625% glycerol were spotted onto an LB-agarose pad and imaged every 2 min at 37 °C. This experiment was done in tandem with Movies S1 and S3.

**Movie S3: Some cells exhibit repeated phases of growth and shrinking during regrowth after overnight passage in LB + 0.625% glycerol, related to Fig. S5A.** Wild-type *B. subtilis* 168 cells from an 18-h culture in LB + 0.625% glycerol were spotted onto an LB-agarose pad and imaged every 2 min at 37 °C. For one cell, no growth was observed for the first 3 h; afterward it engaged in short phases of growth and shrinking, with small blebs forming during growth. Some of its progeny also exhibited periodic shrinking and death through explosive lysis. This experiment was done in tandem with Movies S1 and S2.

## Supplementary Tables

**Table S1:**
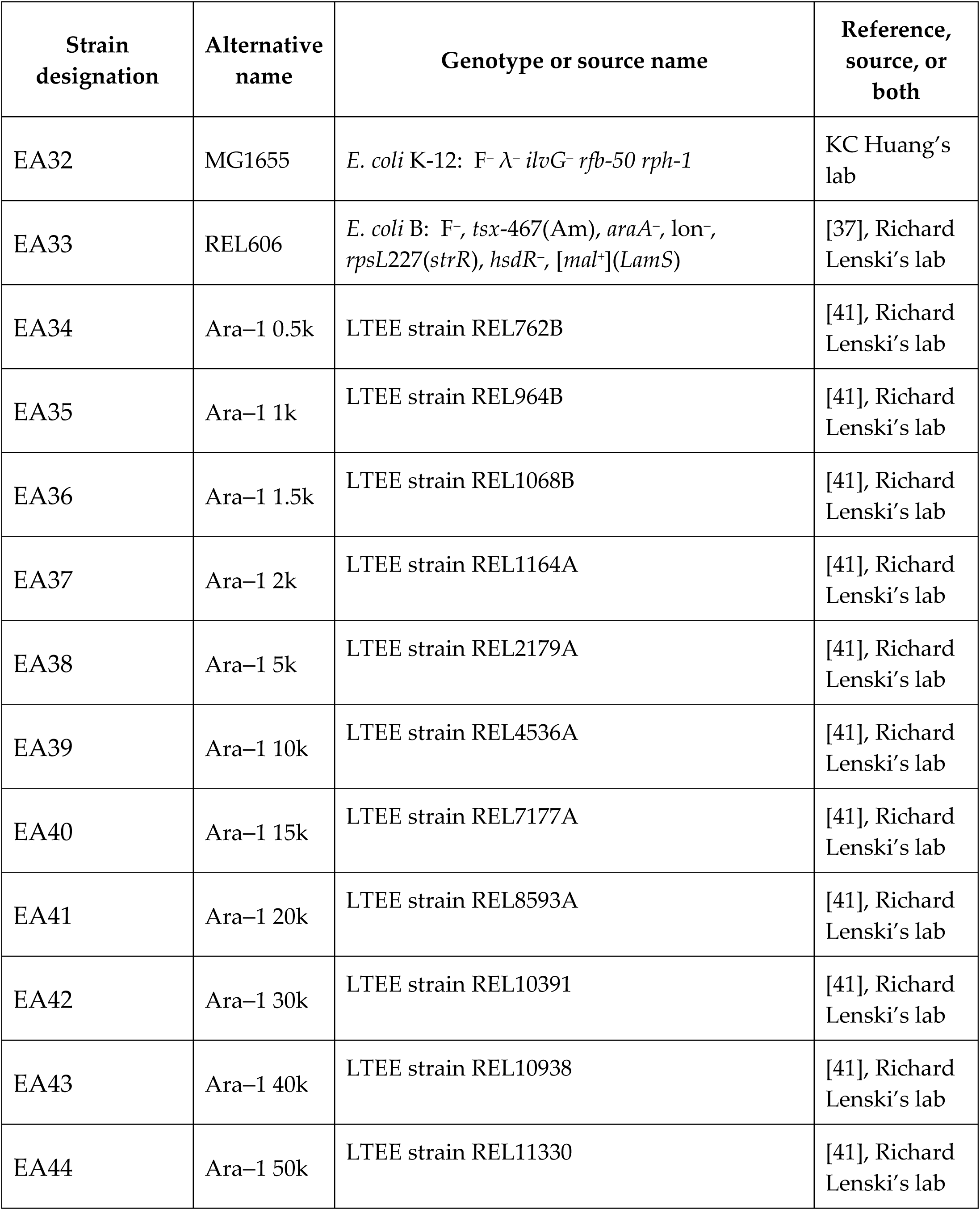
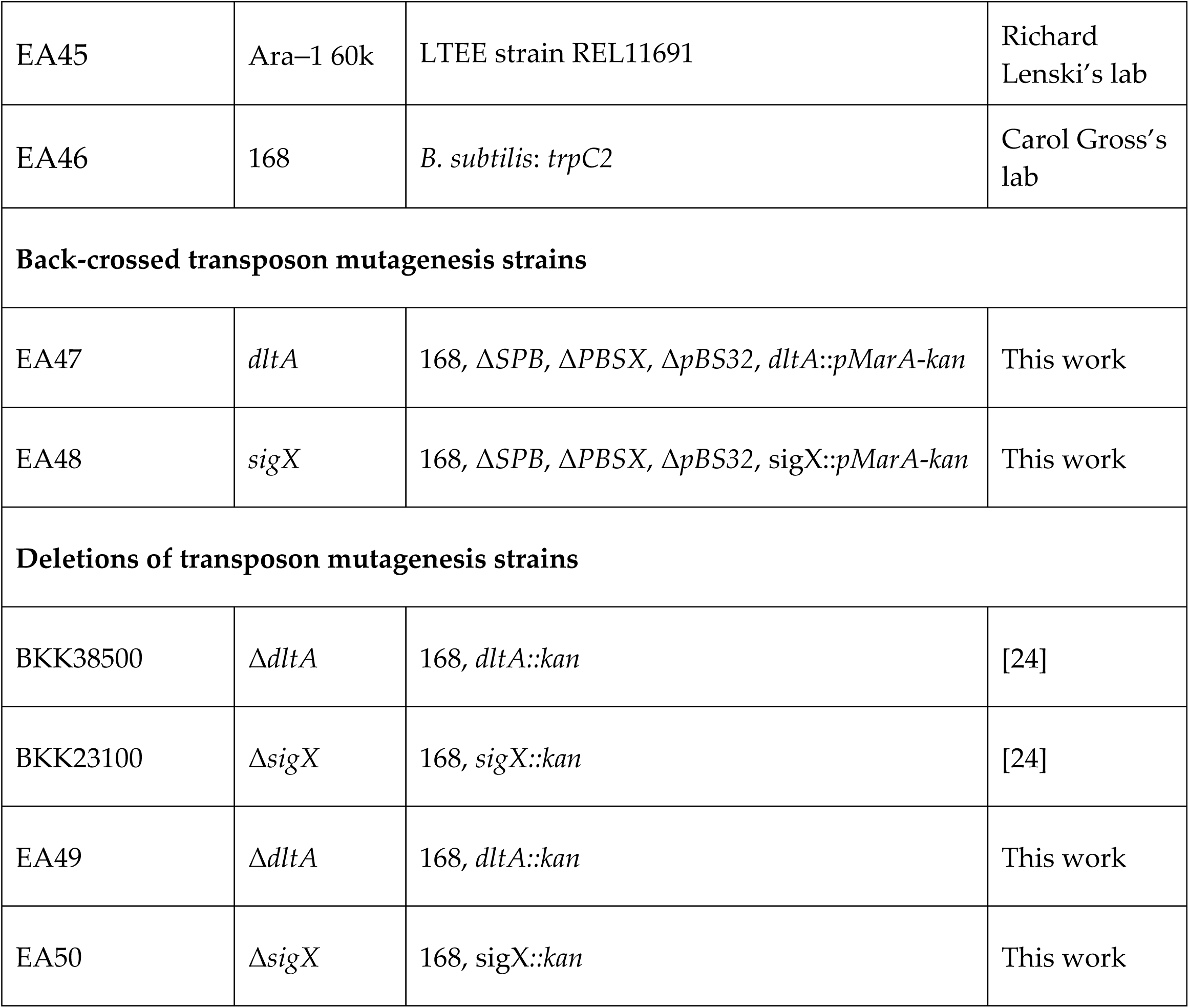
List of strains used in this study.

| Strain designation | Alternative name | Genotype or source name | Reference, source, or both |
| --- | --- | --- | --- |
| EA32 | MG1655 | *E. coli* K-12: F<sup>-</sup> λ-*ilvG*<sup>-</sup> *rfb*-50 *rph*-1 | KC Huang's lab |
| EA33 | REL606 | *E. coli* B: F<sup>-</sup>, *tsx*-467(Am), *araA*<sup>-</sup>, *lon*<sup>-</sup>, *rpsL*227(*strR*), *hsdR*<sup>-</sup>, [*mal*<sup>+</sup>](*LamS*) | [37], Richard Lenski's lab |
| EA34 | Ara-1 0.5k | LTEE strain REL762B | [41], Richard Lenski's lab |
| EA35 | Ara-1 1k | LTEE strain REL964B | [41], Richard Lenski's lab |
| EA36 | Ara-1 1.5k | LTEE strain REL1068B | [41], Richard Lenski's lab |
| EA37 | Ara-1 2k | LTEE strain REL1164A | [41], Richard Lenski's lab |
| EA38 | Ara-1 5k | LTEE strain REL2179A | [41], Richard Lenski's lab |
| EA39 | Ara-1 10k | LTEE strain REL4536A | [41], Richard Lenski's lab |
| EA40 | Ara-1 15k | LTEE strain REL7177A | [41], Richard Lenski's lab |
| EA41 | Ara-1 20k | LTEE strain REL8593A | [41], Richard Lenski's lab |
| EA42 | Ara-1 30k | LTEE strain REL10391 | [41], Richard Lenski's lab |
| EA43 | Ara-1 40k | LTEE strain REL10938 | [41], Richard Lenski's lab |
| EA44 | Ara-1 50k | LTEE strain REL11330 | [41], Richard Lenski's lab |
| EA45 | Ara-1 60k | LTEE strain REL11691 | Richard Lenski's lab |
| EA46 | 168 | *B. subtilis: trpC2* | Carol Gross's lab |
| **Back-crossed transposon mutagenesis strains** |  |  |  |
| EA47 | *dltA* | 168, ΔSPB, ΔPBSX, ΔpBS32, *dltA::pMarA-kan* | This work |
| EA48 | *sigX* | 168, ΔSPB, ΔPBSX, ΔpBS32, *sigX::pMarA-kan* | This work |
| **Deletions of transposon mutagenesis strains** |  |  |  |
| BKK38500 | Δ*dltA* | 168, *dltA::kan* | [24] |
| BKK23100 | Δ*sigX* | 168, *sigX::kan* | [24] |
| EA49 | Δ*dltA* | 168, *dltA::kan* | This work |
| EA50 | Δ*sigX* | 168, *sigX::kan* | This work |

## References

1. Weinstock, M.T., et al., Vibrio natriegens as a fast-growing host for molecular biology. Nat Methods, 2016. 13(10): p. 849–51.

2. Jørgensen, B.B. and S. Hondt, A Starving Majority Deep Beneath the Seafloor. Science, 2006. 314(5801): p. 932.

3. Roszak, D.B. and R.R. Colwell, Survival strategies of bacteria in the natural environment. Microbiological reviews, 1987. 51(3): p. 365–379.

4. Monds, R.D., et al., Systematic perturbation of cytoskeletal function reveals a linear scaling relationship between cell geometry and fitness. Cell Rep, 2014. 9(4): p. 1528–37.

5. Peters, J.M., et al., A Comprehensive, CRISPR-based Functional Analysis of Essential Genes in Bacteria. Cell, 2016. 165(6): p. 1493–1506.

6. Vasi, F., M. Travisano, and R.E. Lenski, Long-term experimental evolution in Escherichia coli. II. Changes in life-history traits during adaptation to a seasonal environment. The american naturalist, 1994. 144(3): p. 432–456.

7. Monod, J., The Growth of Bacterial Cultures. Annual Review of Microbiology, 1949. 3: p. 371–394.

8. Arjes, H.A., et al., Biosurfactant-Mediated Membrane Depolarization Maintains Viability during Oxygen Depletion in Bacillus subtilis. Current Biology, 2020. 30(6): p. 1011-1022.e6.

9. Arjes, Heidi A., et al., Failsafe Mechanisms Couple Division and DNA Replication in Bacteria. Current Biology, 2014. 24(18): p. 2149–2155.

10. Koch, A.L., Some calculations on the turbidity of mitochondria and bacteria. Biochimica et Biophysica Acta, 1961. 51(3): p. 429–441.

11. Koch, A.L., Turbidity measurements of bacterial cultures in some available commercial instruments. Analytical Biochemistry, 1970. 38(1): p. 252–259.

12. Zwietering, M.H., et al., Modeling of the Bacterial Growth Curve. Applied and Environmental Microbiology, 1990. 56(6): p. 1875–1881.

13. Tonner, P.D., et al., Detecting differential growth of microbial populations with Gaussian process regression. Genome research, 2017. 27(2): p. 320–333.

14. Baranyi, J. and T.A. Roberts, A dynamic approach to predicting bacterial growth in food. International Journal of Food Microbiology, 1994. 23(3): p. 277–294.

15. Gibson, A.M., N. Bratchell, and T.A. Roberts, Predicting microbial growth: growth responses of salmonellae in a laboratory medium as affected by pH, sodium chloride and storage temperature. International Journal of Food Microbiology, 1988. 6(2): p. 155–178.

16. Hills, B.P. and K.M. Wright, A New Model for Bacterial Growth in Heterogeneous Systems. Journal of Theoretical Biology, 1994. 168(1): p. 31–41.

17. Jones, J.E., et al., Mathematical modelling of the growth, survival and death of Yersinia enterocolitica. International Journal of Food Microbiology, 1994. 23(3): p. 433–447.

18. Whiting, R.C. and J.E. Call, Time of growth model for proteolytic Clostridium botulinum. Food Microbiology, 1993. 10(4): p. 295–301.

19. Whiting, R.C. and M. Cygnarowicz-Provost, A quantitative model for bacterial growth and decline. Food Microbiology, 1992. 9(4): p. 269–277.

20. Schepers, A.W., J. Thibault, and C. Lacroix, Comparison of simple neural networks and nonlinear regression models for descriptive modeling of Lactobacillus helveticus growth in pH-controlled batch cultures. Enzyme and Microbial Technology, 2000. 26(5): p. 431–445.

21. McKellar, R.C., A heterogeneous population model for the analysis of bacterial growth kinetics. International Journal of Food Microbiology, 1997. 36(2): p. 179–186.

22. Blount, Z.D., et al., Genomic analysis of a key innovation in an experimental Escherichia coli population. Nature, 2012. 489(7417): p. 513–8.

23. Baba, T., et al., Construction of Escherichia coli K-12 in-frame, single-gene knockout mutants: the Keio collection. Mol Syst Biol, 2006. 2: p. 2006 0008.

24. Koo, B.M., et al., Construction and Analysis of Two Genome-Scale Deletion Libraries for Bacillus subtilis. Cell Syst, 2017. 4(3): p. 291–305 e7.

25. Campos, M., et al., Genomewide phenotypic analysis of growth, cell morphogenesis, and cell cycle events in Escherichia coli. Mol Syst Biol, 2018. 14(6): p. e7573.

26. Auer, G.K., et al., Mechanical Genomics Identifies Diverse Modulators of Bacterial Cell Stiffness. Cell Syst, 2016. 2(6): p. 402–11.

27. Trivedi, R.R., et al., Mechanical Genomic Studies Reveal the Role of Alanine Metabolism in Pseudomonas aeruginosa Cell Stiffness. mBio, 2018. 9.

28. Liu, H., et al., Magic Pools: Parallel Assessment of Transposon Delivery Vectors in Bacteria. mSystems, 2018. 3(1).

29. Shiver, A.L., et al., Rapid ordering of barcoded transposon insertion libraries of anaerobic bacteria. bioRxiv, 2019.

30. Liu, H., et al., Large-scale chemical-genetics of the human gut bacterium Bacteroides thetaiotaomicron. bioRxiv, 2019.

31. Colavin, A., H. Shi, and K.C. Huang, RodZ modulates geometric localization of the bacterial actin MreB to regulate cell shape. Nat Commun, 2018. 9(1): p. 1280.

32. Schmidt, A., et al., The quantitative and condition-dependent Escherichia coli proteome. Nat Biotechnol, 2016. 34(1): p. 104–10.

33. Gompertz, B., On the Nature of the Function Expressive of the Law of Human Mortality, and on a New Mode of Determining the Value of Life Contingencies. Philosophical Transactions of the Royal Society of London, 1825. 115: p. 513–583.

34. Shi, H., et al., Deep Phenotypic Mapping of Bacterial Cytoskeletal Mutants Reveals Physiological Robustness to Cell Size. Curr Biol, 2017. 27(22): p. 3419–3429 e4.

35. Feltham, R.K.A., et al., A Simple Method for Storage of Bacteria at −76°C. Journal of Applied Bacteriology, 1978. 44(2): p. 313–316.

36. Perry, S.E., Freeze-Drying and Cryopreservation of Bacteria in Methods in Molecular Biology: Cryopreservaflon and Freeze-Drying Protocols J.G. Day and M.R. Mclellan, Editors. 1995, Humana Press Inc.: Totowa, NJ p. 21–30.

37. Lenski, R.E., et al., Long-Term Experimental Evolution in Escherichia coli. I. Adaptation and Divergence During 2,000 Generations. The American Naturalist, 1991. 138(6): p. 1315–1341.

38. Studier, F.W., et al., Understanding the differences between genome sequences of Escherichia coli B strains REL606 and BL21(DE3) and comparison of the E. coli B and K-12 genomes. J Mol Biol, 2009. 394(4): p. 653–80.

39. Wiser, M.J., N. Ribeck, and R.E. Lenski, Long-term dynamics of adaptation in asexual populations. Science, 2013. 342(6164): p. 1364–7.

40. Good, B.H., et al., The dynamics of molecular evolution over 60,000 generations. Nature, 2017. 551(7678): p. 45–50.

41. Tenaillon, O., et al., Tempo and mode of genome evolution in a 50,000-generation experiment. Nature, 2016. 536(7615): p. 165–70.

42. Novak, M., et al., Experimental tests for an evolutionary trade-off between growth rate and yield in E. coli. Am Nat, 2006. 168(2): p. 242–51.

43. Singh, K.D., et al., Carbon catabolite repression in Bacillus subtilis: quantitative analysis of repression exerted by different carbon sources. J Bacteriol, 2008. 190(21): p. 7275–84.

44. Petit, M.A., et al., Tn10-derived transposons active in Bacillus subtilis. Journal of bacteriology, 1990. 172(12): p. 6736–6740.

45. Cao, M. and J.D. Helmann, The Bacillus subtilis extracytoplasmic-function sigmaX factor regulates modification of the cell envelope and resistance to cationic antimicrobial peptides. J Bacteriol, 2004. 186(4): p. 1136–46.

46. Heaton, M.P. and F.C. Neuhaus, Biosynthesis of D-alanyl-lipoteichoic acid: cloning, nucleotide sequence, and expression of the Lactobacillus casei gene for the D-alanine-activating enzyme. Journal of Bacteriology, 1992. 174(14): p. 4707.

47. Neuhaus, F.C. and J. Baddiley, A continuum of anionic charge: structures and functions of D-alanyl-teichoic acids in gram-positive bacteria. Microbiol Mol Biol Rev, 2003. 67(4): p. 686–723.

48. Wormann, M.E., et al., Enzymatic activities and functional interdependencies of Bacillus subtilis lipoteichoic acid synthesis enzymes. Mol Microbiol, 2011. 79(3): p. 566–83.

49. Parker, R.F., Action of penicillin on Staphylococcus; effect of size of inoculum on the test for sensitivity. Proceedings of the Society for Experimental Biology and Medicine. Society for Experimental Biology and Medicine (New York, N.Y.), 1946. 63(2): p. 443–446.

50. Stevenson, K., et al., General calibration of microbial growth in microplate readers. Sci Rep, 2016. 6: p. 38828.

51. Aranda-Diaz, A., et al., Bacterial interspecies interactions modulate pH-mediated antibiotic tolerance. Elife, 2020. 9.

52. Blount, Z.D., C.Z. Borland, and R.E. Lenski, Historical contingency and the evolution of a key innovation in an experimental population of Escherichia coli. Proc Natl Acad Sci U S A, 2008. 105(23): p. 7899–906.

53. Grosskopf, T., et al., Metabolic modelling in a dynamic evolutionary framework predicts adaptive diversification of bacteria in a long-term evolution experiment. BMC Evol Biol, 2016. 16(1): p. 163.

54. Maddamsetti, R., R.E. Lenski, and J.E. Barrick, Adaptation, Clonal Interference, and Frequency-Dependent Interactions in a Long-Term Evolution Experiment with Escherichia coli. Genetics, 2015. 200(2): p. 619–31.

55. Rozen, D.E. and R.E. Lenski, Long-Term Experimental Evolution in Escherichia coli. VIII. Dynamics of a Balanced Polymorphism. Am Nat, 2000. 155(1): p. 24–35.

56. Lenski, R.E. and M. Travisano, Dynamics of adaptation and diversification: a 10,000-generation experiment with bacterial populations. Proceedings of the National Academy of Sciences, 1994. 91(15): p. 6808.

57. Schaechter, M., O. MaalØe, and N.O. Kjeldgaard, Dependency on Medium and Temperature of Cell Size and Chemical Composition during Balanced Growth of Salmonella typhimurium. Microbiology, 1958. 19(3): p. 592–606.

58. Mongold, J.A. and R.E. Lenski, Experimental rejection of a nonadaptive explanation for increased cell size in Escherichia coli. J Bacteriol, 1996. 178(17): p. 5333–4.

59. Schirner, K., et al., Distinct and essential morphogenic functions for wall- and lipo-teichoic acids in Bacillus subtilis. The EMBO journal, 2009. 28(7): p. 830–842.

60. Carlton, B.C. and B.J. Brown, Gene mutation, in Manual of methods for general bacteriolog, P. Gerhardt, Editor. 1981, American Society for Microbiology: Washington, D.C. p. 222–242

61. Edelstein, A., et al., Computer control of microscopes using microManager. Curr Protoc Mol Biol, 2010. Chapter 14: p. Unit14 20.

62. Yasbin, R.E. and F.E. Young, Transduction in Bacillus subtilis by Bacteriophage SPP1. Journal of Virology, 1974. 14(6): p. 1343.

63. Miller, J.H., A Short Course in Bacterial Genetics: A Laboratory Manual and Handbook for Escherichia Coli and Related Bacteria. 1992: Cold Spring Harbor Laboratory Press.

64. Sambrook, J., Molecular cloning: a laboratory manual, ed. D.W. Russell. 2001, Cold Spring Harbor, N.Y: Cold Spring Harbor Laboratory.

65. Le Breton, Y., N.P. Mohapatra, and W.G. Haldenwang, In vivo random mutagenesis of Bacillus subtilis by use of TnYLB-1, a mariner-based transposon. Applied and Environmental Microbiology, 2006. 72(1): p. 327–333.

